# DORA AI Scientist: Multi-agent Virtual Research Team for Scientific Exploration Discovery and Automated Report Generation

**DOI:** 10.1101/2025.03.06.641840

**Authors:** Vladimir Naumov, Diana Zagirova, Sha Lin, Yupeng Xie, Wenhao Gou, Anatoly Urban, Nina Tikhonova, Khadija Alawi, Mike Durymanov, Fedor Galkin, Shan Chen, Denis Sidorenko, Mike Korzinkin, Morten Scheibye-Knudsen, Alan Aspuru-Guzik, Evgeny Izumchenko, David Gennert, Frank W. Pun, Man Zhang, Petrina Kamya, Alex Aliper, Feng Ren, Alex Zhavoronkov

## Abstract

Modern goal-oriented scientific research process involves hierarchical teams of researchers of diverse backgrounds performing generalist and domain-specific tasks. Many of these tasks include hypothesis generation, literature review, data collection, cleanup, processing and analysis, experimental design, virtual and physical experiments, research report and academic paper writing, reference management, bibliography and quality control. Most of these tasks can be performed automatically or in a co-pilot mode by the generative reinforcement learning systems. In this paper, we introduce a versatile multi-agent scientific exploration and draft outline research assistant (DORA), which provides multiple templates and workflows for automated or semi-automated research studies and report generation. Under user guidance, it employs hierarchical teams of AI agents based on the plug-and-play generalist and domain-specific large language models (LLMs) exploiting a variety of specialized research tools and open data repositories and generates high-quality research outputs publication drafts with maximally-accurate references. DORA is designed to minimize the time and effort required for manuscript preparation, thereby enabling researchers to devote more attention to high-value discovery tasks. The system is constantly evolving with user feedback with regular feature and resource updates. The platform is available at https://dora.insilico.com.

## Introduction

Scientific writing is an essential component of the research process, serving not only as a vehicle for disseminating findings but also as a foundation for peer review and the advancement of interdisciplinary knowledge. In today’s fast-paced environment, marked by rapid technological progress and innovation, there is a growing need to refine and streamline the writing process. The traditional approach to writing is inherently complex, involving a range of demanding tasks such as data curation, comprehensive literature reviews, grammatical precision, linguistic clarity, stylistic enhancement, and meticulous citation management. Scientific writing often requires collaborative efforts from entire research teams, leading to substantial investments of time and resources. These challenges can significantly impede researchers’ ability to convey their findings effectively, ultimately hindering the advancement of knowledge and innovation [1], [2]. As a result, scientific writing emerges as a bottleneck, delaying the timely dissemination of critical insights within the scientific community. In light of these challenges, there is a clear demand for innovative technologies that can facilitate and enhance the scientific writing process.

Recent advances in large language models (LLMs) such as those developed by Anthropic, Google DeepMind, and OpenAI, have opened new avenues for tackling challenges in scientific writing and enhancing the speed of knowledge dissemination through AI assistance [3], [4], [5], [6], [7]. These models, including ChatGPT, have demonstrated remarkable abilities to process large amounts of data, generate insights, and assist researchers across various disciplines [8]. Their ability to produce coherent, human-like text from minimal inputs makes LLMs particularly well-suited for the creation of AI writing assistants. Recent introductions of AI writers, predominantly based on popular open-source LLMs, have attempted automation tasks such as literature searches, summarizing research findings, and drafting manuscript sections [9]. These AI tools range from general-purpose applications to those specifically designed for academic writing [10], [11], [12], [13] and enhancing literature search methodologies [14], [15]. Despite their potential, these tools face significant limitations due to the inherent constraints of LLMs. A major concern is the quality of the information produced, which often lacks the depth and nuance required for sophisticated academic writing. AI tools frequently struggle with complex scientific concepts and technical terminology, resulting in inaccuracies and superficial content [16]. Moreover, these tools are often unable to integrate diverse data sources or perform domain-specific tasks critical for scientific writing. High rates of hallucination—where the AI generates plausible yet incorrect information—along with misleading references, further compromise the reliability of AI-generated content. Additionally, these tools generally lack the capability to conduct original research or synthesize novel ideas, which limits their effectiveness in academic contexts.

These limitations could be particularly mitigated with the advent of autonomous LLM agents. Autonomous LLM agents are AI systems that integrate advanced natural language processing capabilities with the ability to use external tools and execute multi-step tasks [17]. Unlike traditional LLMs, these agents generate human-like text and interact with external sources to perform complex sequences of actions. Agent systems improve performance for diverse applications, especially those requiring complex task solving. The initial applications of these agents span from personal assistants to coding, showcasing their versatility and potential [18]. LLM agents have been further augmented by the development of multi-agent systems, which facilitate communication between individual agents, thereby enhancing their collective capabilities and imitate human step-by-step solution performing. This evolution has been supported by appearance intuitive agents frameworks such as LangChain and LlamaIndex [19], [20], as well as various multi-agent frameworks [21], [22], [23], [24]. These frameworks have significantly boosted the deployment and functionality of autonomous agents across various domains. One of the pioneering applications of autonomous LLM agents in research was the introduction of “AI scientists,” which demonstrated the potential of these agents to function as independent research systems [25]. Subsequent studies have highlighted the promising role of these agents in other domain areas [26], [27]. Going even more deeper, one of the most recent pipelines introduced an agent-based framework for the fully autonomous conducting computational science research with further drafting of academic texts [28]. Thus, LLM agents seem to open a significant opportunity to address the limitations of current AI academic writers through their ability to autonomously carry out complex, multi-step goals and establish a connection without various data resources.

In this paper, we introduce Science42: DORA (Draft Outline Research Assistant), an advanced AI-driven multi-agent tool aimed at improving the scientific drafting process (Fig. 1). DORA is part of a broader suite of AI solutions developed by Insilico Medicine. Since the emergence of large language models (LLMs), Insilico Medicine has swiftly incorporated these technologies into its framework, offering various LLM-based tools, including a biomedical assistant that employs a knowledge graph and a chat-based model for knowledge acquisition across three primary AI platforms focused on target discovery (Fig. 2).

**Fig 1:**
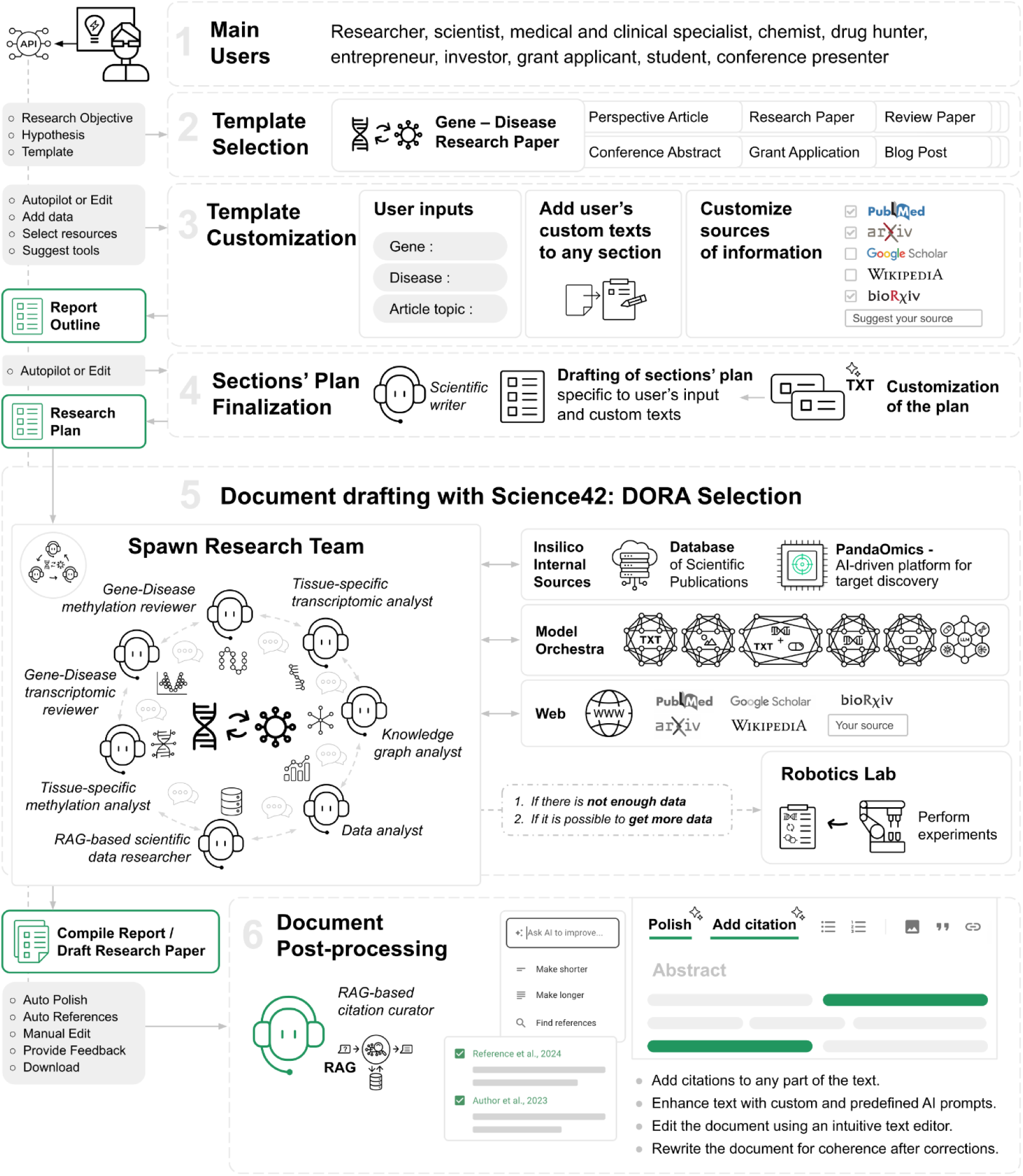
Generalized multi-agent scientific exploration and report generation workflow facilitated by DORA

**Fig. 2.**
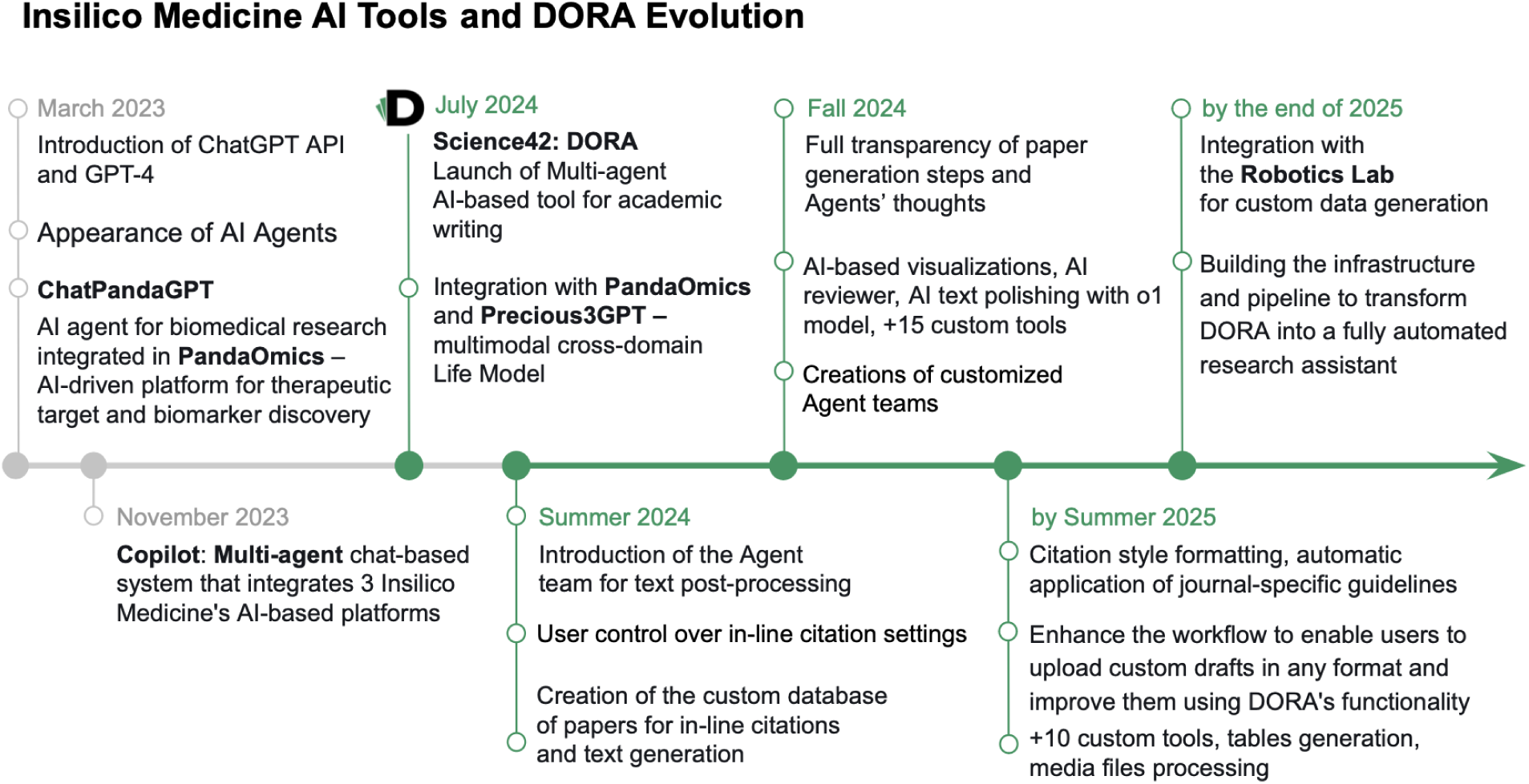
Timeline of AI-Driven Insilico Medicine Products Emergence and DORA development

DORA utilizes a multi-agent system, enhanced with custom tools linked to a variety of data sources, to systematically assist in the creation of effective scientific texts. This platform synthesizes knowledge from Insilico Medicine’s internal resources, proprietary models, and external databases, ultimately converting this information into publishable content for the scientific community. Its integration with internal platforms significantly enhances its capabilities, particularly in the biomedical domain and the area of target discovery. DORA addresses the limitations of existing AI writing tools by automating the drafting and search for scientific references, allowing researchers to focus on their primary investigative tasks rather than the time-consuming writing process. Additionally, DORA has the potential to develop into an independent research assistant as it uses a chain-of-thought approach that underpins its multi-agent system. This approach is seen as a promising direction for future AI systems, possibly giving them reasoning abilities that allow them to perform tasks on their own and generate new ideas [29]. Our research aims to demonstrate DORA’s functionalities and highlight its potential to transform the scientific writing landscape.

### DORA pipeline overview

**Step 1-2 - template selection.** DORA is engineered to streamline the drafting process, enabling users to prepare document drafts with minimal effort. The process initiates with the selection of a template from a curated collection, which includes a variety of document types, ranging from research papers focused on drug discovery to general blog posts (Fig. 1). Each template functions as a sophisticated pipeline that manages the entire document generation workflow. Specifically, each template is a JSON file that defines the overall composition of the final document, designated data sources for content generation, formatting and style guidelines, and meticulously crafted prompts based on the latest best practices, among other settings. Importantly, these templates are not static; they are flexible frameworks that can be tailored to address the specific research requirements of the user. Prior to the generation process, the selected JSON template can be customized through the user interface, ensuring that even a single template can yield unique outputs based on user-specific adjustments. The reliance on predefined templates allows seamless draft creation without requiring users to have prior experience with LLMs or interact with complex prompts, while maintaining flexibility by allowing user-defined inputs, predefined text segments, or modifications to section outlines. DORA offers over 20 predefined templates designed for various purposes, including research articles, review papers, perspective pieces, cover letters, blog posts, press releases, grant applications, patent submissions, medical case studies, and more.

**Step 3 - template customization.** Upon selecting the most appropriate template, the customization process initiates by defining specific inputs, which may include the research topic, gene, or disease under investigation. These inputs are aligned with the parameters of the chosen template. The interface subsequently offers suggestions for potential input variations. One of the key customization steps is the ability to provide predefined text for each section. DORA’s agents use this information throughout the generation process, ensuring a smooth combination of predefined and generated content. For example, if the user has already drafted parts of the Results section, this text can be inserted into the corresponding section in the interface. This data is then used to create the remaining parts of the Results section and inform the generation of other related sections, such as the Introduction and Abstract.

DORA supports the selection of information sources for paper generation used by agents in the context generation phase. We accommodate a wide range of sources suitable for various purposes, including scientific databases such as PubMed, ArXiv, and Google Scholar, as well as broader sources like Wikipedia and Google Search. Furthermore, we allow users to upload their own documents to serve as primary sources of information and citations during the drafting process.

**Step 4 - outline drafting.** The generation of complex sections, which require sophisticated data analysis and comprehension, is executed in two steps: initially, a plan is created, followed by the construction of the section itself. Specialized LLM agents create these plans, ensuring they are comprehensive and specific to the user’s defined inputs or custom texts. Users have the flexibility to modify these plans according to their specific requirements, including the addition, alteration, or removal of topics. Adjusting the outline allows to shift the overall focus of the section or fine-tune specific subtopics to be emphasized in the final text.

**Step 5 - document generation.** Upon customization, DORA orchestrates the research process, delivering a high-quality publication draft. This is achieved through the deployment of multiple LLM agents, each assigned specific roles focusing on different sections of the manuscript. These agents leverage LLM enhanced with custom tools, enabling the retrieval of both Insilico Medicine’s internal data and external databases via API calls. Supplementary Table 1 presents the primary tool groups currently available in DORA. Each agent operates under precisely engineered prompts aligned with the chosen template and the specific section being generated. For initialization of the agents, we utilize LLM library that comprises local and cloud-based solutions. We employ the LangChain [20] framework but make a range of custom modifications.

The current agents are tightly integrated within the Insilico Medicine AI-based infrastructure though the set of custom tools. Specifically, agents have access to PandaOmics, our AI-driven, cloud-based platform designed for target discovery [30], as well as proprietary models such as Precious3GPT [31]. Precious3GPT is a multi-modal, multi-omics, multi-species language model designed to represent omics changes and predict targets and compounds that can reverse those changes. We plan to further enhance this integration. One of the crucial aspects of this initiative is the interconnection with the Robotics Lab established by Insilico Medicine, where autonomous guided robotics conduct a series of laboratory experiments [32]. This integration aims to facilitate data generation, thereby enriching the dataset on the biological subjects that form the primary focus of the current publication initiated by the user. This represents one of the steps towards developing DORA into a full-stack autonomous research assistant.

**Step 6.1 - text post-processing: citation finder.** The generated article appears section by section in the WYSIWYG text editor. Further interaction with LLM agents can continue on the initial draft to improve text quality or customize it further. One of the key features is a citation finder. Any text segment can be highlighted to activate the agent with the Retrieval-Augmented Generation (RAG) system, which searches for the most suitable citations. The user receives a list of available citations to choose from, and the bibliography on the right side of the page updates accordingly.

**Step 6.2 - text post-processing: AI actions**. Additionally, any text part could be highlighted to activate the LLM agents that could increase or reduce the size of the text part or perform any action with the text based on the user’s defined request. These agents are empowered with the background knowledge of the articles used in the paper generation, ensuring high-quality text reduction or expansion without introducing vacuity into the final paper.

**Step 6.3 - text post-processing: polish text**. As we provide features for post-generation text rewriting, we created an additional feature “Polish” that makes the entire document consistent in terms of the content. If any facts were removed or added, the corresponding texts of the changed section and all dependent sections (refer to “Data interconnection and memory flow section” for dependencies description) would be regenerated. This feature ensures that making a coherent document even after extensive manual correction would take minimal manual effort.

While the LLM agents adhere to the research workflow defined by the template, leveraging pre-defined research resources and databases, the collaborative effort of agents working in a step-by-step manner ensures the creation of a unique research product. This results in not only a polished and coherent document but also valuable insights that can drive broader scientific exploration. It makes DORA not just a tool for generating research publications, but a catalyst for innovation, empowering researchers to push the boundaries of scientific discovery by interaction with a human in the loop.

### Detailed methodology and key pipeline features

#### Beyond a Conventional AI Writer

DORA is engineered not merely to generate text on a given topic but to elucidate research trajectories and shape them into coherent, structured narratives. Distinguishing itself from other AI writing assistants, DORA leverages specialized agents connected with our proprietary research platforms and publication databases. Such integration allows DORA to gather the knowledge from diverse sources and then combine them to get a novel conclusion.

Among other specialized agents with integration to other platforms, we introduced a type integrated with our extensive internal publication database. This agent leverages Retrieval-Augmented Generation (RAG) to perform thorough data collection and analysis, significantly reducing the risk of hallucinations and ensuring transparency by providing relevant links to scientific publications. RAG operates by extracting information from a curated database of over 28 million scientific texts from PubMed and PMC. The implementation of RAG is enhanced by an additional layer that formulates multiple queries from the initial question, thereby enhancing the quality of the content and reducing the susceptibility to syntax formulation errors. Besides, we rely on the hybrid mode of the retrieval that combines full-text search by keywords and vector search with further merging and re-ranking of chunks. This implementation was found to be optimal in capturing both direct inclusion of the main words and semantic similarity. RAG agents are responsible for comprehensive data collection and citation insertion during text generation, ensuring accuracy and depth. Usage of RAG agent during the generation is one of the ways to reduce the hallucinations rates of LLMs. As provided with the wide range of data and instructed to mainly rely on it in the scientific data discussions, ensures that the generated content is valid and based on reliable scientific information.

#### Integration with Insilico Medicine platforms

The composition of agents within each section and their available tools is matched to the document type and focus. Domain or topic-specific templates are empowered with agents which are able to retrieve information from AI-based platforms of Insilico Medicine. For instance, a Gene-Disease research paper leverages domain-specific LLM agents integrated with PandaOmics, an AI-driven platform aimed to facilitate automatic target discovery [30]. These agents contain tools to access an extensive database of processed omics datasets, curated knowledge graphs, and other validated internal sources useful for biomedical drafts preparation. While DORA’s flexibility allows for the incorporation of predefined results, it can also autonomously guide research without preliminary data independently gathering and synthesizing information from PandaOmics. The integration with PandaOmics enables DORA to efficiently extract processed data, substantiate a gene’s involvement in a disease without relying on pre-existing results and construct a reliable molecular hypothesis on gene involvement in the disease onset. The integration with domain-specific platforms not just facilitates the creation of well-structured, insightful publications but also enriches research by offering valuable insights for future scientific exploration.

One groundbreaking integration within DORA is the incorporation of Insilico Medicine’s cutting-edge Precious3GPT model for aging research [31]. We integrated into one of the DORA’s agents that gained an ability to effectively guide the prediction of prospective geroprotectors. The agents work collaboratively to generate a research article draft, detailing the promising geroprotectors identified by P3GPT based on user-specified inputs.

#### Data interconnection and memory flow

Among solutions to enhance the quality of individual sections, DORA also offers features that elevate the entire research pipeline. One of DORA’s standout features is a shared memory system that interlinks all output sections, enhancing the logical flow and coherence of the final document. It implies the initiation of generation with sections that involve data collection and analysis, such as the Results in a research article. Once completed, these texts are stored in the memory of LLM agents working on the article. If the user provides the custom texts, they are also stored in the memory and used for further generations. Only after these data-centric sections are completed, the LLM agents begin creating descriptive sections like the Introduction and Abstract. While generating these sections, the LLM agents are instructed to refer to the stored memory to incorporate insights from previous iterations. This pipeline guarantees that major results, insights, and conclusions are explicitly discussed across all interconnected sections, resulting in well-structured and cohesive publications. For this feature, we employed the ConversationSummaryBufferMemory functionality from LangChain, which combines two key components. First, it maintains a buffer of recent interactions (or sections, in our case) in memory. When the buffer reaches its capacity, instead of discarding older interactions entirely, it compiles them into a summary that is also utilized. Given our reliance on the long-context LLM model, we opted to retain a substantial number of tokens in the buffer to minimize information loss.

We observed that, in some cases, the LLM agent is not capable of extracting precise details from the memory. However, the preservation of details is exceptionally important for the construction of dependent sections. To address this limitation, we introduced a second type of dependency: the insertion of the previous section directly into the prompt of the dependent section. To keep token usage lower, this approach was employed only in cases where the preservation of details from previous iterations is crucial.

Thus, each template establishes two types of dependencies: through the agent’s memory and through the direct insertion of ready sections into the prompt of a generated one. The final setting of a section within a template usually follows this structure:

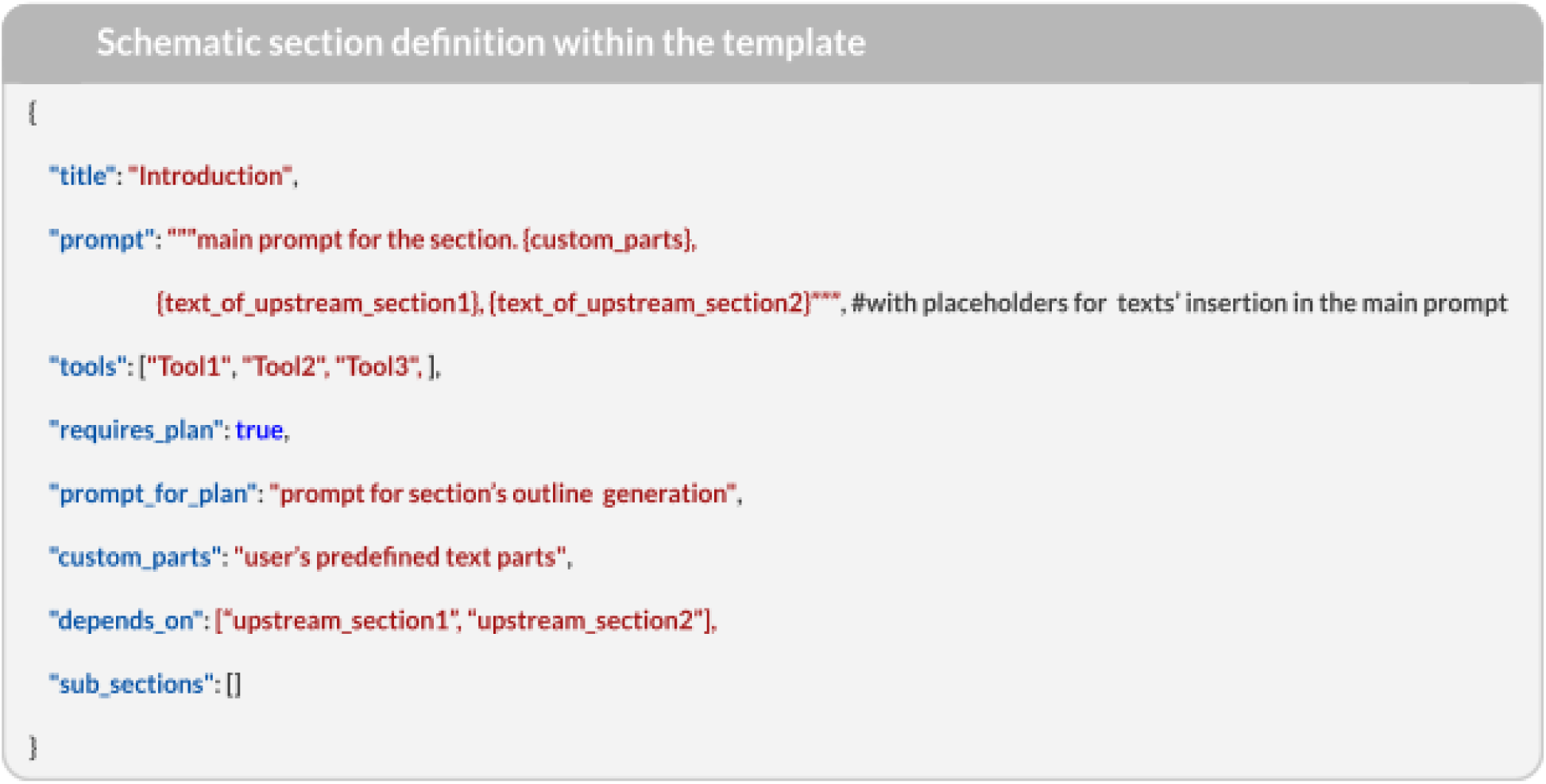

These two dependencies are considered to finalize the order of section generation. Specifically, sections with no dependencies are generated first, while the order of other sections is defined based on the representation of the dependencies as a directed graph and the application of a topological sorting algorithm. Thus, the final pipeline logic is unique for each of the templates and documented in the JSON file. The schematic representation of the pipeline with dependencies is shown on Figure 3.

**Fig. 3.**
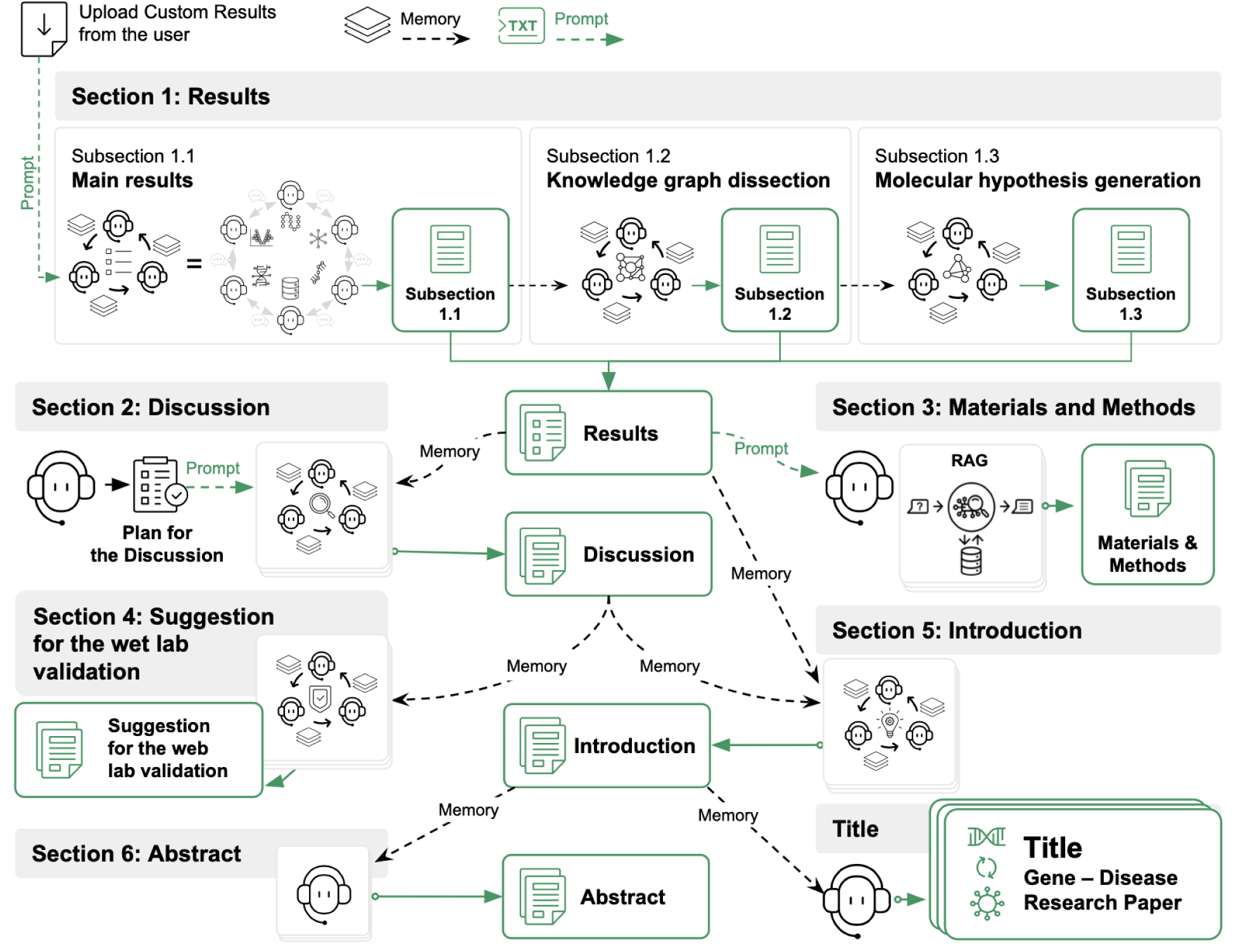
The pipeline logic for the generation of the research paper related to the identification of the selected Gene involvement into the Disease.

**Fig. 4.**
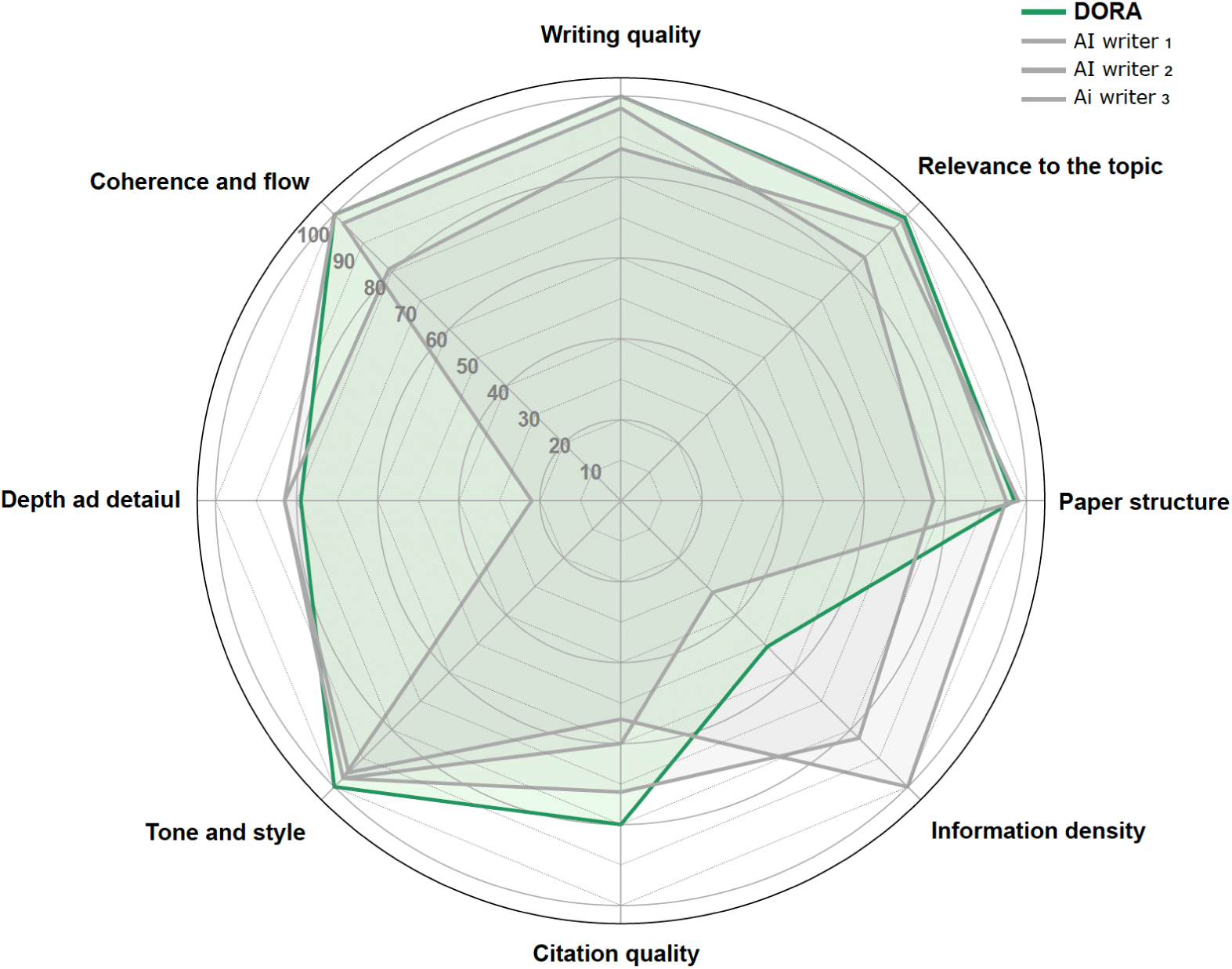
Median normalized scores for papers generated by DORA and 3 selected AI writers.

### Case studies

#### Applications of DORA in Advancing Biomedical Research

● https://www.medrxiv.org/content/10.1101/2024.07.04.24309952v1
● https://www.biorxiv.org/content/10.1101/2024.10.02.616224v1
● https://www.biorxiv.org/content/10.1101/2024.12.02.626364v1
● https://www.biorxiv.org/content/10.1101/2025.01.09.632065v1.full
● https://www.biorxiv.org/content/10.1101/2024.07.25.605062v1.full

## Applications of DORA in Advancing Biomedical Research

The integration of artificial intelligence (AI) in biomedical research is revolutionizing the way scientists analyze complex datasets, generate hypotheses, and draft scientific manuscripts. DORA, an advanced AI writing tool, plays a pivotal role in facilitating the research and publication process by helping researchers structure their findings, refine their arguments, and accelerate the drafting of scientific articles. The diversity of research topics, methodologies, and applications across different areas of biomedical science highlights DORA’s versatility in supporting various research endeavors.

One notable example is the development of **Precious3GPT (P3GPT)**, a multimodal, multi-species, multi-omics, and multi-tissue transformer model for aging research and drug discovery. P3GPT enables tasks such as age prediction across species, target discovery, and drug sensitivity prediction, demonstrating an advanced AI-driven approach to biological data analysis. Similarly, in the study **AI-Driven Toolset for IPF and Aging Research**, researchers explored the aging-related mechanisms of idiopathic pulmonary fibrosis (IPF) using a pathway-aware proteomic aging clock. The findings established novel connections between aging biology and IPF pathogenesis, underscoring the potential of AI-guided therapeutic **development.** In both cases, DORA supported the drafting process by organizing complex research findings into a structured narrative, streamlining the articulation of AI methodologies, and ensuring clarity and coherence in scientific communication.

In the domain of infectious diseases, DORA assisted in drafting the article **Designing a Multi-Serotype Dengue Virus Vaccine: An In Silico Approach to Broad-Spectrum Immunity**. This study leveraged computational tools to design a vaccine capable of eliciting broad immune responses against multiple dengue virus serotypes. The research incorporated in silico methods such as B-cell and T-cell epitope prediction, toxicity screening, and structural validation of the vaccine construct. With DORA’s assistance, researchers efficiently structured their results, providing a clear and comprehensive description of the vaccine design process and its potential impact on global health.

Beyond infectious diseases and aging research, DORA has also contributed to oncology studies, such as the **Comparative Study of Radiotherapy Outcomes Across Cancer Types**. This research analyzed radiotherapy outcomes in glioblastoma multiforme (GBM) and low-grade gliomas (LGG), revealing molecular mechanisms underlying variations in treatment efficacy. By assisting in drafting this article, DORA enabled the authors to present their findings in a logically coherent manner, and provide comprehensive introduction

These examples illustrate the broad applicability of DORA in biomedical research, encompassing different topics, methodologies, and application areas, from ML, drug discovery and aging biology to infectious disease modeling and oncology. By enhancing the efficiency and quality of scientific writing, DORA empowers researchers to focus on innovation and discovery while ensuring their findings are communicated with clarity and precision.

As AI continues to shape the landscape of academic science, tools like Science42: DORA will remain indispensable in accelerating literature review, and translating complex results into structured and accurate content

### Links -

Comparative analysis of Endoxifen, Tamoxifen and Fulvestrant: A Bioinformatics Approach to Uncover Mechanisms of Action in Breast Cancer - [33]

Designing a multi-serotype Dengue virus vaccine: an in silico approach to broad-spectrum immunity - [34]

AI-Driven Toolset for IPF and Aging Research Associates Lung Fibrosis with Accelerated Aging - [35]

Precious3GPT: Multimodal Multi-Species Multi-Omics Multi-Tissue Transformer for Aging Research and Drug Discovery - [31]

https://www.google.com/url?q = https://www.medrxiv.org/content/10.1101/2024.07.04.24309952v1&sa=D&source=editors&ust=1740560203566430&usg=AOvVaw3c_DFpmmfBEc0-mIRriWxR - [36]

DORA has demonstrated its utility in drafting high-impact research articles across diverse fields. Multiple AI-focused publications from Insilico Medicine have benefited from DORA’s comprehensive research capabilities during the drafting process. Notable examples include the Precious3GPT research paper and the study “AI-Driven Toolset for IPF and Aging Research Associates Lung Fibrosis with Accelerated Aging” [31], [35]. In these projects, DORA supported the drafting process by organizing complex research findings into structured narratives, streamlining the retrieval of supporting information, drafting literature reviews, ensuring clarity and coherence, and significantly increasing the efficiency of paper preparation.

Beyond extensive internal applications, DORA has extended its reach to external collaborators, establishing itself as a valuable asset in the broader scientific community. In oncology, DORA facilitated a groundbreaking collaboration with Professor Morten Scheibye-Knudsen at the University of Copenhagen. This partnership resulted in an innovative primary research article examining molecular signatures that influence radiotherapy outcomes in glioblastoma multiforme and low-grade gliomas [36]. Working with Dr. Filippo Castiglione at the Technology Innovation Institute, DORA demonstrated its adaptability in infectious disease research by supporting the development of a manuscript on multi-serotype Dengue virus vaccine design [34]. Most recently, DORA played a crucial role in facilitating research collaboration between Atossa Therapeutics and Insilico Medicine, contributing significantly to a publication on novel approaches to breast cancer treatment [33]. These diverse applications conclusively demonstrate DORA’s exceptional versatility and effectiveness in enhancing scientific research productivity. By facilitating literature review, automatic data search and analysis, application of custom templates and tools, manuscript preparation across multiple biomedical disciplines, DORA stands as an invaluable resource for researchers.

### Examples of the generated articles

DORA is designed to generate a variety of document types across multiple knowledge domains. However, its primary strength lies in producing scientific texts within the biomedical field. There are several examples of such documents and general types of original research articles.

## Example 1: Research article focused on the identification of the mechanisms of the Gene involvement into the Disease onset considering the user’s defined molecular hypothesis

User’s selected inputs:

1. Gene - CDKN3;
2. Disease - Hepatocellular carcinoma;
3. Hypothesis - CDKN3 is a therapeutic target for Hepatocellular carcinoma.

Please note that the parts of the text that were corrected in using citation finder and AI features in DORA, or manually corrected in the in-built text editor, are highlighted in green.

Research paper

#### Therapeutic Targeting of CDKN3: A Promising Strategy for the Treatment of Hepatocellular Carcinoma (HCC)

##### Abstract

Hepatocellular carcinoma (HCC) remains a major global health issue due to its high incidence, mortality rates, and limited treatment options. Current therapies often fail to provide long-term benefits, highlighting the urgent need for novel therapeutic targets. Our study examines CDKN3 (cyclin-dependent kinase inhibitor 3) as a potential therapeutic target for HCC. Using an artificial intelligence-driven approach that combines multiomics data analysis and knowledge graph dissection, we aimed to clarify the role of CDKN3 in HCC pathogenesis. Differential expression analysis revealed marked upregulation of CDKN3 in HCC tissues and activation of CDKN3-associated signaling pathways, suggesting its involvement in cell cycle regulation. Knowledge graph analysis further identified the interaction of CDKN3 with the Akt and other downstream signaling pathways, which are crucial for cell proliferation and survival in HCC. By integrating omics data and advanced pathway analysis, we provided the rationale for targeting CDKN3 as a promising treatment option in HCC. These findings pave the way for the development of CDKN3 inhibitors and their evaluation in preclinical and clinical settings. Our work also demonstrates the importance of integrating advanced bioinformatics tools and multiomics data to uncover critical regulatory networks in cancer that could be used for drug discovery.

##### Introduction

Identifying therapeutic targets is a crucial phase in the drug discovery process across various diseases, playing a vital role in the development of effective treatments. Hepatocellular carcinoma (HCC) is one of the most common and deadly forms of liver cancer, with a high incidence and mortality rate worldwide (Le Grazie et al. 2017). Despite advances in medical research, the prognosis for HCC patients remains poor due to late diagnosis and limited treatment options. Current therapies, including surgical resection, liver transplantation, and systemic treatments like sorafenib, often fail to provide long-term benefits and are associated with high recurrence rates and significant side effects (Tejeda-Maldonado et al. 2015; Tanaka, Shimada, and Kudo 2014). This highlights the urgent need for novel therapeutic targets and strategies to improve patient outcomes.

Recent advancements in AI-driven methodologies have shown promise in identifying potential therapeutic targets, even in complex diseases, by analyzing multiomics data and utilizing advanced pathway analysis (Pun et al. 2023; Visan and Negut 2024). These innovative approaches can address the limitations of traditional target identification methods, which often fail to account for the complexity and heterogeneity of diseases like HCC.

In this context, our study focuses on CDKN3 (cyclin-dependent kinase inhibitor 3) as a potential therapeutic target for HCC. CDKN3 is overexpressed in various cancers, including HCC, and is implicated in tumor progression and chemotherapeutic resistance. CDKN3 promotes tumor growth by facilitating the G1-S transition in the cell cycle and is associated with poor prognosis in cancer patients (Mori et al. 2022). Despite its potential significance, the specific mechanisms by which CDKN3 contributes to HCC pathology remain poorly understood. Addressing this challenge, this study aimed to evaluate CDKN3 as a therapeutic target for HCC by integrating omics data, network analysis, and literature evidence. The performed analysis is based on data from PandaOmics, a cloud-based platform that leverages artificial intelligence and bioinformatics to process multimodal omics and biomedical text data, aiming to identify therapeutic targets and biomarkers (Kamya et al. 2024).

Our findings support the hypothesis that therapeutic targeting of CDKN3 is a promising strategy for the treatment of HCC. The overexpression of CDKN3, its role in activating the Akt signaling pathway, and its interaction with key cell cycle regulators highlight its potential as a crucial therapeutic target.

##### Materials and Methods

###### PandaOmics Platform

PandaOmics, developed by Insilico Medicine, is a cloud-based software that uses artificial intelligence and bioinformatics to analyze multimodal omics and biomedical text data for identifying therapeutic targets and biomarkers. It provides a detailed data processing pipeline, combining gene expression, proteomics, and methylation data, and uses meta-analyses to improve target predictions. The platform aids therapeutic target discovery through an intuitive interface, utilizing 23 disease-specific models and incorporating biological knowledge graphs and large language models for precise analysis.

###### Gene Expression Analysis

Gene expression datasets related to HCC patients (GSE113409, GSE77269, GSE113392, GSE136319, GSE67170, GSE83691, GDC-TCGA-LIHC, TCGA-LIHC, GSE45267, GSE67764, GSE59259, GSE19665, E-MTAB-5905, GSE82177, GSE60502, GSE6764, GSE102079, GSE107170, and GSE36376) were processed using PandaOmics. The platform automatically detects data types and suggests normalization methods. Upper-quartile normalization and log2-transformation were applied to GEO datasets, while only log2-transformation was used for UCSC Xena cancer datasets. Differential analysis was performed using the limma package, with p-values adjusted by the Benjamini–Hochberg procedure. Meta-analysis in PandaOmics calculated combined logarithmic fold-changes (LogFC) and Q-values across datasets using minmax normalization and Stouffer’s method for p-value combination, followed by false discovery rate (FDR) correction.

###### Methylation Analysis

Methylation data for HCC was also processed using PandaOmics. The platform handled the methylation datasets (GSE67170, GSE113392, GSE113409, TCGA-LIHC, GSE83691, GSE136319, GSE77269) by detecting data types and suggesting suitable normalization methods. Differential methylation analysis was conducted using the limma package, with p-values adjusted by the Benjamini–Hochberg procedure. Combined logarithmic fold-changes (LogFC) and Q-values were calculated across datasets using minmax normalization and Stouffer’s method for p-value combination, followed by FDR correction.

###### Pathway activation analysis

The analysis of altered pathways was conducted in PandaOmics using the iPANDA algorithm (Ozerov et al. 2016) to compare HCC tumors versus normal tissue in patients, utilizing data from GSE113409, GSE77269, GSE113392, GSE136319, GSE67170, GSE83691, GDC-TCGA-LIHC, TCGA-LIHC, GSE45267, GSE67764, GSE59259, GSE19665, E-MTAB-5905, GSE82177, GSE60502, GSE6764, GSE102079, GSE107170, and GSE36376 datasets. The Reactome signaling pathway graph served as the database for the iPANDA algorithm (Jassal et al. 2020). This approach determines the direction and intensity of pathway activation by applying a linear combination of logarithmic fold-changes, along with statistical and topological weights assigned to each gene within the pathway, as described in (Ozerov et al. 2016). Consequently, the iPANDA algorithm output illustrates how gene expression variations between case and control groups influence signaling pathway levels, with each pathway’s score referred to as the iPANDA score. High iPANDA scores indicate upregulation of pathways, whereas low scores suggest downregulation. Pathway perturbations are visualized on a pathway diagram, with each node color-coded according to the average logarithmic fold-change values of the genes within that node.

###### Knowledge Graph Analysis

PandaOmics assists researchers in target and biomarker selection and provides strong evidence to support each hypothesis. By integrating bulk multi-omics data and scientific literature insights through an advanced biological knowledge graph, the platform offers detailed information on genes, diseases, biological processes, and compounds. It also uses clinical trial data to provide a deeper understanding of the competitive landscape.

###### AI-driven paper writing tool

Writing of the initial manuscript draft was performed using DORA, Insilico Medicine LLM-based paper drafter assistant, which generates templated scientific paper drafts. The process of paper generation is curated by multiple AI agents, powered by Large Language Models (LLMs), and integrated with PandaOmics internal databases. Each agent employs Retrieval-Augmented Generation (RAG) to perform comprehensive data collection and analysis, to reduce the probability of hallucinations, and to provide relevant PubMed links to make the generation of the paper more transparent. The final version of the manuscript was thoroughly curated manually.

##### Results

###### Methylation and expression analysis of the *CDKN3* gene

To investigate the role of CDKN3 in HCC, we utilized the PandaOmics platform, which integrates artificial intelligence and bioinformatics techniques to analyze multimodal omics and biomedical text data for therapeutic target and biomarker discovery.

The combined differential expression analysis showed that CDKN3 is significantly upregulated in HCC as compared to normal tissues, with a normalized log fold change (LogFC) of 0.769 and a highly significant p-value of 7.97e-144. Further examination of individual comparisons highlighted that CDKN3 is consistently upregulated across all examined HCC datasets, including GSE36376, GDC-TCGA-LIHC, GSE102079, and eight others, with LogFC values ranging from 0.31 to 0.953, suggesting a strong association with the disease state.

In contrast, the methylation data indicates a slight downregulation of CDKN3 in HCC, with a combined LogFC of -0.195 and a p-value of 1.58e-08. This downregulation is observed across several datasets, including TCGA-LIHC and GSE77269, with LogFC values of -0.116 and -0.095, respectively. The discrepancy between expression and methylation data suggests that while CDKN3 expression is upregulated, its promoter region may be hypomethylated, potentially contributing to its overexpression. The consistency of these findings across different datasets highlights the potential regulatory role of methylation in CDKN3 expression.

###### Signaling pathway analysis

To further understand the role of CDKN3 in HCC, we performed Pathway Activation Network Decomposition Analysis (iPANDA). This scalable and robust method allows to identify activated signaling pathways using gene expression data. The CDKN3 gene is associated with 11 signaling pathways in the Reactome integrated platform for biological pathways. It turned out that 10 of 11 pathways exhibited positive iPANDA scores with highest values for “mitotic G1 and G1-S transition”, “mitotic G2-G2-M phases”, and “regulation of mitotic cell cycle” across the majority of individual comparisons. This suggests a strong association between increased CDKN3 expression and high cell proliferation rate in HCC.

###### Knowledge graph dissection

The knowledge graph analysis provided additional insights into the biological connections between CDKN3 and HCC. First, CDKN3 is associated with the Akt signaling pathway, which is crucial for cell survival, proliferation, and response to chemotherapy in HCC and other cancers (Dai et al. 2016; Yu et al. 2020; Chen et al. 2024). In our analysis, expression levels of *AKT1* show slight alterations, with LogFC values ranging from -0.088 to -0.146. Interaction of CDKN3 with microRNAs adds another layer of regulation for this pathway. These non-coding RNAs modulate CDKN3 expression and activity, influencing the downstream signaling pathways and contributing to the malignant phenotype of HCC. For instance, miR-181d-5p-mediated inactivation of the Akt signaling pathway through CDKN3 suppression could lead to reduced cancer cell proliferation and invasion as was shown in non-small cell lung cancer cells (Gao et al. 2019). CDKN3 also forms complexes with Ki67 and p53 proteins (Zargar et al. 2024), downstream targets of the Akt pathway, which are essential for cell proliferation and HCC development. MKI67, a marker for cell proliferation, shows significant upregulation across almost all HCC datasets with a combined LogFC of 0.569 and a p-value of 3.4e-96, while *TP53* shows insignificant alteration, with LogFC values ranging from -0.193 to 0.063.

Additionally, the BIRC5 gene has been identified by a functional network analysis as the central node in the CDKN3 co-expression network in HCC, displaying the highest number of interactions with other genes (Xing et al. 2012). BIRC5, also known as survivin, is part of the inhibitor of apoptosis (IAP) gene family, which encodes proteins that inhibit apoptotic cell death (Salvesen and Duckett 2002). In terms of cell proliferation, BIRC5 acts as a component of the chromosomal passenger complex, aiding in chromosome segregation and cytokinesis (Giodini et al. 2002). Additionally, BIRC5 may enhance the nuclear accumulation of cyclin D and CDK4, leading to increased phosphorylation of Rb and promoting G1/S phase progression (Connell, Wheatley, and McNeish 2008). Our results show that BIRC5 is significantly upregulated in HCC. The combined differential expression analysis revealed a combined LogFC of 0.55 with a highly significant p-value of 5.77e-103, while the methylation analysis of BIRC5 in HCC showed a slight downregulation with a combined LogFC of -0.099 and a p-value of 0.0005. These findings offer deeper insights into the molecular mechanisms through which CDKN3 influences cell cycling in HCC.

Next, CDKN3 interacts with the *MYCN* gene, a well-known oncogene, and other cell cycle proteins such as CDC6 and CDK4, forming a network that promotes their mutual expression in neuroblastoma cells (Vernaza et al. 2024). Pathways such as the cell cycle regulation pathway, involving CDKN3, CDC6, and CDK4, and the differentiation pathways modulated by *MYCN*, are crucial in the context of HCC pathology as well (Yasukawa et al. 2020). All the mentioned CDKN3 interactors display significant upregulation across almost all HCC datasets with combined LogFC values of 0.588, 0.418, and 0.2 for *CDC6*, *CDK4*, and *MYCN*, respectively.

Finally, the cell cycle regulation pathway, where CDKN3 interacts with phosphatase KAP, is crucial for maintaining proper cell cycle checkpoints (Okamoto, Kitabayashi, and Taya 2006; Hannon, Casso, and Beach 1994), which is often disrupted in HCC (Yeh et al. 2000).

#### Discussion

Our study aimed to evaluate CDKN3 as a therapeutic target for hepatocellular carcinoma by integrating omics data, network analysis, and literature evidence.

Analysis of the differential expression data for CDKN3 indicates its robust overexpression and hypomethylation in HCC tissues compared to normal liver tissues that is consistent with experimental data in HCC tumors and cell lines (Xing et al. 2012; Mori et al. 2022). Noteworthy, this overexpression is associated with poor prognosis for HCC (Dai et al. 2020), and other cancers (Chang et al. 2018; Vernaza et al. 2024; Li et al. 2014; Wang et al. 2019; Fan et al. 2015; Al Sharie et al. 2023).

Network analysis has shown that CDKN3 (cyclin-dependent kinase inhibitor 3) plays a significant role in hepatocellular carcinoma pathology. Although CDKN3 is known to regulate cell cycle progression via dephosphorylation of CDK2 kinase (Nalepa et al. 2013), it is often dysregulated in various cancers, including HCC. Presumably, CDKN3 promotes tumor progression by accelerating the G1-S transition in the cell cycle and is associated with poor prognosis in cancer patients (Mori et al. 2022). Consistent with these data, among the genes associated with cell cycle regulation, genomic studies have identified the CDKN3 as a gene with abnormal expression and well-established diagnostic and prognostic values for HCV-associated HCC (Zhang et al. 2021).

Based on our findings, we hypothesized that the Akt signaling pathway appears to be a crucial link between CDKN3 and HCC, with CDKN3 modulating the activity of Akt1 and its downstream targets including Ki67, p53, and BIRC5 (Gao et al. 2019) resulting in altered cell survival and proliferation. According to our data, expression levels of AKT1 and TP53 exhibit minor fluctuations in HCC. However, experimental inhibition of CDKN3 leads to altered expression of p53, p21, and Akt, suggesting its role in tumor differentiation and chemotherapeutic tolerance (Dai et al. 2016). It should be noted that downregulation of CDC6 and CDK4 after CDKN3 inhibition (Vernaza et al. 2024) could also be mediated by affecting the Akt signaling pathway, which activates cyclin D1/Cdk4 signaling (Shimura et al. 2012).

To confirm our hypothesis, further experimental studies should be conducted to determine the mechanism of the CDKN3/Akt interaction. To date, CDK2 is the only known substrate of CDKN3 phosphatase (Nalepa et al. 2013). However, CDKN3 may have other substrates, which could directly or indirectly contribute to activation of Akt. For example, dephosphorylation of phosphatases that negatively regulate Akt, like PP2A and PTEN, or dephosphorylation of kinases that activate Akt, like Src, could be putative mechanisms of the CDKN3/Akt interplay.

Although CDKN3 phosphatase plays an important role in pathogenesis of multiple cancers, no specific pharmacological inhibitors have been reported so far. At the same time, development of small-molecule inhibitors of CDKN3 seems to be a feasible strategy because this phosphatase belongs to a druggable protein family. Alternatively, the RNAi approach could be exploited for CDKN3 targeting. The hypomethylation of the CDKN3 promoter in HCC tumors indicates an epigenetic mechanism that increases CDKN3 expression, which also could be targeted for therapeutic intervention.

In summary, our analysis supports the hypothesis that therapeutic targeting of CDKN3 is a promising strategy for the treatment of hepatocellular carcinoma. The overexpression of CDKN3, its role in activating the Akt signaling pathway, and its interaction with key cell cycle regulators highlight its potential as a critical therapeutic target. Further research should focus on the mechanisms of the CDKN3/Akt interplay and the development of CDKN3 inhibitors, along with evaluation of their efficacy in preclinical and clinical settings to improve outcomes for HCC patients.

References

Al Sharie, Ahmed H., Abdulmalek M. Abu Zahra, Tamam El-Elimat, Reem F. Darweesh, Ayah K. Al-Khaldi, Balqis M. Abu Mousa, Mohammad S. Bani Amer, et al. 2023. “Cyclin Dependent Kinase Inhibitor 3 (CDKN3) Upregulation Is Associated with Unfavorable Prognosis in Clear Cell Renal Cell Carcinoma and Shapes Tumor Immune Microenvironment: A Bioinformatics Analysis.” *Medicine* 102 (36): e35004.

Chang, Shih-Lun, Tzu-Ju Chen, Ying-En Lee, Sung-Wei Lee, Li-Ching Lin, and Hong-Lin He. 2018. “CDKN3 Expression Is an Independent Prognostic Factor and Associated with Advanced Tumor Stage in Nasopharyngeal Carcinoma.” *International Journal of Medical Sciences* 15 (10): 992–98.

Chen, Yingjun, Dai Li, Kaihui Sha, Xuezhong Zhang, and Tonggang Liu. 2024. “Human Pan-Cancer Analysis of the Predictive Biomarker for the CDKN3.” *European Journal of Medical Research* 29 (1): 272.

Connell, Claire M., Sally P. Wheatley, and Iain A. McNeish. 2008. “Nuclear Survivin Abrogates Multiple Cell Cycle Checkpoints and Enhances Viral Oncolysis.” *Cancer Research* 68 (19): 7923–31.

Dai, Wei, Shuo Fang, Guanhe Cai, Jialiang Dai, Guotai Lin, Qiurong Ye, Huilai Miao, et al. 2020. “CDKN3 Expression Predicates Poor Prognosis and Regulates Adriamycin Sensitivity in Hepatocellular Carcinoma in Vitro.” *The Journal of International Medical Research* 48 (7): 300060520936879.

Dai, Wei, Huilai Miao, Shuo Fang, Tao Fang, Nianping Chen, and Mingyi Li. 2016. “CDKN3 Expression Is Negatively Associated with Pathological Tumor Stage and CDKN3 Inhibition Promotes Cell Survival in Hepatocellular Carcinoma.” *Molecular Medicine Reports* 14 (2): 1509–14.

Fan, Chao, Lu Chen, Qingling Huang, Tao Shen, Eric A. Welsh, Jamie K. Teer, Jianfeng Cai, W. Douglas Cress, and Jie Wu. 2015. “Overexpression of Major CDKN3 Transcripts Is Associated with Poor Survival in Lung Adenocarcinoma.” *British Journal of Cancer* 113 (12): 1735–43.

Gao, Li-Ming, Yue Zheng, Ping Wang, Lei Zheng, Wen-Li Zhang, Ya Di, Lan-Lan Chen, et al. 2019. “Tumor-Suppressive Effects of microRNA-181d-5p on Non-Small-Cell Lung Cancer through the CDKN3-Mediated Akt Signaling Pathway in Vivo and in Vitro.” *American Journal of Physiology. Lung Cellular and Molecular Physiology* 316 (5): L918–33.

Giodini, Alessandra, Marko J. Kallio, Nathan R. Wall, Gary J. Gorbsky, Simona Tognin, Pier Carlo Marchisio, Marc Symons, and Dario C. Altieri. 2002. “Regulation of Microtubule Stability and Mitotic Progression by Survivin.” *Cancer Research* 62 (9): 2462–67.

Hannon, G. J., D. Casso, and D. Beach. 1994. “KAP: A Dual Specificity Phosphatase That Interacts with Cyclin-Dependent Kinases.” *Proceedings of the National Academy of Sciences of the United States of America* 91 (5): 1731–35.

Jassal, Bijay, Lisa Matthews, Guilherme Viteri, Chuqiao Gong, Pascual Lorente, Antonio Fabregat, Konstantinos Sidiropoulos, et al. 2020. “The Reactome Pathway Knowledgebase.” *Nucleic Acids Research* 48 (D1): D498–503.

Kamya, Petrina, Ivan V. Ozerov, Frank W. Pun, Kyle Tretina, Tatyana Fokina, Shan Chen, Vladimir Naumov, et al. 2024. “PandaOmics: An AI-Driven Platform for Therapeutic Target and Biomarker Discovery.” *Journal of Chemical Information and Modeling* 64 (10): 3961–69.

Le Grazie, Marco, Maria Rosa Biagini, Mirko Tarocchi, Simone Polvani, and Andrea Galli. 2017. “Chemotherapy for Hepatocellular Carcinoma: The Present and the Future.” *World Journal of Hepatology* 9 (21): 907–20.

Li, T., H. Xue, Y. Guo, and K. Guo. 2014. “CDKN3 Is an Independent Prognostic Factor and Promotes Ovarian Carcinoma Cell Proliferation in Ovarian Cancer.” *Oncology Reports*. https://www.spandidos-publications.com/or/31/4/1825.

Mori, Jinichi, Takahiro Sawada, Taisuke Baba, Akira Hayakawa, Yoshiaki Kanemoto, Koichi Nishimura, Rei Amano, et al. 2022. “Identification of Cell Cycle-Associated and -Unassociated Regulators for Expression of a Hepatocellular Carcinoma Oncogene Cyclin-Dependent Kinase Inhibitor 3.” *Biochemical and Biophysical Research Communications* 625 (October):46–52.

Nalepa, Grzegorz, Jill Barnholtz-Sloan, Rikki Enzor, Dilip Dey, Ying He, Jeff R. Gehlhausen, Amalia S. Lehmann, et al. 2013. “The Tumor Suppressor CDKN3 Controls Mitosis.” *The Journal of Cell Biology* 201 (7): 997–1012.

Okamoto, Koji, Issay Kitabayashi, and Yoichi Taya. 2006. “KAP1 Dictates p53 Response Induced by Chemotherapeutic agents via Mdm2 Interaction.” *Biochemical and Biophysical Research Communications* 351 (1): 216–22.

Ozerov, Ivan V., Ksenia V. Lezhnina, Evgeny Izumchenko, Artem V. Artemov, Sergey Medintsev, Quentin Vanhaelen, Alexander Aliper, et al. 2016. “In Silico Pathway Activation Network Decomposition Analysis (iPANDA) as a Method for Biomarker Development.” *Nature Communications* 7 (November):13427.

Pun, Frank W., Geoffrey Ho Duen Leung, Hoi Wing Leung, Jared Rice, Tomas Schmauck-Medina, Sofie Lautrup, Xi Long, et al. 2023. “A Comprehensive AI-Driven Analysis of Large-Scale Omic Datasets Reveals Novel Dual-Purpose Targets for the Treatment of Cancer and Aging.” *Aging Cell* 22 (12): e14017.

Salvesen, Guy S., and Colin S. Duckett. 2002. “IAP Proteins: Blocking the Road to Death’s Door.” *Nature Reviews. Molecular Cell Biology* 3 (6): 401–10.

Shimura, T., N. Noma, T. Oikawa, Y. Ochiai, S. Kakuda, Y. Kuwahara, Y. Takai, A. Takahashi, and M. Fukumoto. 2012. “Activation of the AKT/cyclin D1/Cdk4 Survival Signaling Pathway in Radioresistant Cancer Stem Cells.” *Oncogenesis* 1 (6): e12.

Tanaka, Katsuaki, Mitsuo Shimada, and Masatoshi Kudo. 2014. “Characteristics of Long-Term Survivors Following Sorafenib Treatment for Advanced Hepatocellular Carcinoma: Report of a Workshop at the 50th Annual Meeting of the Liver Cancer Study Group of Japan.” *Oncology* 87 Suppl 1 (November):104–9.

Tejeda-Maldonado, Javier, Ignacio García-Juárez, Jonathan Aguirre-Valadez, Adrián González-Aguirre, Mario Vilatobá-Chapa, Alejandra Armengol-Alonso, Francisco Escobar-Penagos, Aldo Torre, Juan Francisco Sánchez-Ávila, and Diego Luis Carrillo-Pérez. 2015. “Diagnosis and Treatment of Hepatocellular Carcinoma: An Update.” *World Journal of Hepatology* 7 (3): 362–76.

Vernaza, Alexandra, Daniela F. Cardus, Jadyn L. Smith, Veronica Partridge, Amy L. Baker, Emma G. Lewis, Angela Zhang, Zhenze Zhao, and Liqin Du. 2024. “Identification of CDKN3 as a Key Gene That Regulates Neuroblastoma Cell Differentiation.” *Journal of Cancer* 15 (5): 1153–68.

Visan, Anita Ioana, and Irina Negut. 2024. “Integrating Artificial Intelligence for Drug Discovery in the Context of Revolutionizing Drug Delivery.” *Life* 14 (2). https://doi.org/10.3390/life14020233.

Wang, J., W. Che, W. Wang, G. Su, and T. Zhen. 2019. “CDKN3 Promotes Tumor Progression and Confers Cisplatin Resistance via RAD51 in Esophageal Cancer.” *Cancer Management and Research*. https://www.tandfonline.com/doi/abs/10.2147/CMAR.S193793.

Xing, Chunyang, Haiyang Xie, Lin Zhou, Wuhua Zhou, Wu Zhang, Songming Ding, Bajin Wei, Xiaobo Yu, Rong Su, and Shusen Zheng. 2012. “Cyclin-Dependent Kinase Inhibitor 3 Is Overexpressed in Hepatocellular Carcinoma and Promotes Tumor Cell Proliferation.” *Biochemical and Biophysical Research Communications* 420 (1): 29–35.

Yasukawa, Ken, Lee Chuen Liew, Keitaro Hagiwara, Ai Hironaka-Mitsuhashi, Xian-Yang Qin, Yutaka Furutani, Yasuhito Tanaka, et al. 2020. “MicroRNA-493-5p-Mediated Repression of the MYCN Oncogene Inhibits Hepatic Cancer Cell Growth and Invasion.” *Cancer Science* 111 (3): 869–80.

Yeh, C. T., S. C. Lu, T. C. Chen, C. Y. Peng, and Y. F. Liaw. 2000. “Aberrant Transcripts of the Cyclin-Dependent Kinase-Associated Protein Phosphatase in Hepatocellular Carcinoma.” *Cancer Research*. https://aacrjournals.org/cancerres/article-abstract/60/17/4697/506503.

Yu, Hanxu, Jun Yao, Mingyu Du, Jinjun Ye, Xia He, and Li Yin. 2020. “CDKN3 Promotes Cell Proliferation, Invasion and Migration by Activating the AKT Signaling Pathway in Esophageal Squamous Cell Carcinoma.” *Oncology Letters* 19 (1): 542–48.

Zargar, Seema, Tanveer A. Wani, Salman Alamery, and Fatimah Yaseen. 2024. “Olmutinib Reverses Thioacetamide-Induced Cell Cycle Gene Alterations in Mice Liver and Kidney Tissues, While Wheat Germ Treatment Exhibits Limited Efficacy at Gene Level.” *Medicina* 60 (4). https://doi.org/10.3390/medicina60040639.

Zhang, Yongqiang, Yuqin Tang, Chengbin Guo, and Gen Li. 2021. “Integrative Analysis Identifies Key mRNA Biomarkers for Diagnosis, Prognosis, and Therapeutic Targets of HCV-Associated Hepatocellular Carcinoma.” *Aging* 13 (9): 12865–95.

## Example 2: Review paper

User’s selected topic: Biomarkers in amyotrophic lateral sclerosis: diagnostic and prognostic significance

Review paper

#### Biomarkers in Amyotrophic Lateral Sclerosis: Diagnostic and Prognostic Significance

##### Abstract

Amyotrophic lateral sclerosis (ALS) is a devastating neurodegenerative disorder characterized by the progressive loss of motor neurons, leading to muscle weakness and eventual paralysis. The identification of reliable biomarkers for ALS is critical for early diagnosis, monitoring disease progression, and evaluating therapeutic responses. This review aims to provide a comprehensive overview of the current state of biomarker research in ALS, highlighting the types and significance of various biomarkers, including genetic, biochemical, neurophysiological, cerebrospinal fluid (CSF), and imaging data. We discuss the potential of blood biomarkers for differential diagnosis and prognostic assessment, the role of neurofilaments and other protein markers, and the emerging significance of metabolic and immune markers. Additionally, we explore the challenges and limitations in the validation and clinical application of these biomarkers. The review underscores the urgent need for continued research to develop and validate biomarkers that can be integrated into clinical practice and therapeutic development. Future directions include the standardization of biomarker discovery methodologies, the incorporation of biomarkers into clinical trials, and the exploration of novel biomarkers that reflect the multifaceted nature of ALS. By advancing our understanding and application of ALS biomarkers, we can improve patient outcomes and accelerate the development of effective treatments.

##### Introduction

Despite extensive research, the pathogenesis of ALS remains poorly understood, and there is currently no cure. The identification of reliable biomarkers is critical for early diagnosis, monitoring disease progression, and evaluating therapeutic responses. Biomarkers can provide valuable insights into the underlying mechanisms of ALS and facilitate the development of targeted therapies <u>(Costa Júlia & de Carvalho Mamede, 2016</u>).

The search for ALS biomarkers has intensified in recent years, driven by the need to improve diagnostic accuracy and patient outcomes. Blood biomarkers, in particular, have shown promise in providing a more direct basis for differential diagnosis and prognostic assessment beyond clinical manifestations and electromyography findings (<u>Lv Yongting & Li</u> <u>Hongfu, 2024</u>). However, despite the discovery of numerous candidate biomarkers, none have yet been validated as reliable tools for clinical use or drug development <u>(Benatar</u> <u>Michael</u> et al., <u>2016</u>).

Current limitations in ALS biomarker research include variability in biomarker levels among patients, insufficient sensitivity and specificity, and a lack of longitudinal studies tracking biomarker changes over the course of the disease (Sturmey Ellie & Malaspina Andrea, 2022; Benatar Michael et al., 2016). These challenges underscore the need for more comprehensive research to improve the reliability and utility of ALS biomarkers.

Recent advancements in computational tools and databases have shown potential in addressing these limitations. For example, platforms like Perseus and Gorilla have been developed to analyze proteomics data and identify enriched Gene Ontology terms, respectively, aiding researchers in uncovering new biomarkers for ALS <u>(Bhadra Pratiti</u> et al., <u>2021</u>). These innovative approaches are helping to enhance our understanding of ALS and pave the way for the development of more effective diagnostic and therapeutic strategies.

One of the most pressing challenges in ALS research is the identification of biomarkers that can be applied to clinical practice and incorporated into the development of innovative therapies (Sanchez-Tejerina Daniel et al., 2023). Objective markers of disease activity are highly coveted to stratify the heterogeneous ALS phenotype and facilitate the discovery of effective disease-modifying treatments (Barritt Andrew W et al., 2018). The development of brain imaging biomarkers, for instance, is essential for advancing diagnosis, stratification, and monitoring of ALS in both clinical practice and trials (Mazón Miguel et al., 2018).

This review aims to provide a comprehensive overview of the current state of biomarker research in ALS, highlighting recent advancements, current limitations, and future directions. By synthesizing the latest findings, we hope to offer insights that will guide future research efforts and ultimately improve patient care and outcomes.

#### Biomarkers in Amyotrophic Lateral Sclerosis: Types and Significance

ALS is a rapidly progressive neurodegenerative disease that affects upper and lower motor neurons, with no effective treatment despite numerous clinical trials. Efforts have been made to identify novel disease biomarkers to support diagnosis, provide prognostic information, measure disease progression in trials, and increase knowledge of disease pathogenesis <u>(Costa Júlia & de Carvalho Mamede, 2016</u>). Biomarkers have become a focus of intense research in ALS, with the hope that they might aid therapy development efforts. Despite the discovery of many candidate biomarkers, none have yet emerged as validated tools for drug development (<u>Benatar Michael</u> et al., <u>2016</u>).

Blood biomarkers in ALS can provide a more direct basis for differential diagnosis and prognostic assessment beyond clinical manifestations and electromyography findings (Lv Yongting & Li Hongfu, 2024). Specific biomarkers could help in early detection and diagnosis, and could also act as indicators of disease progression and therapy effectiveness (Takahashi Ikuko et al., 2015). The discovery and validation of biomarkers that can be applied to clinical practice and incorporated into the development of innovative therapies is one of the most explored areas of research in ALS (Sanchez-Tejerina Daniel et al., 2023).

A study was undertaken to determine the diagnostic and prognostic value of a panel of serum biomarkers and to correlate their concentrations with several clinical parameters in a large cohort of ALS patients (Falzone Yuri Matteo et al., 2022). Utilizing neurodegenerative markers for the diagnostic evaluation of ALS is crucial as a definitive diagnostic test or biomarker for ALS is currently unavailable, leading to a diagnostic delay following the onset of initial symptoms (Klíčová Kateřina et al., 2024).

Objective markers of disease sensitive to clinical activity, symptomatic progression, and underlying substrates of neurodegeneration are highly coveted in ALS to stratify the highly heterogeneous phenotype and facilitate the discovery of effective disease-modifying treatments (Barritt Andrew W et al., 2018). Multiple candidate biomarkers for ALS have emerged across a range of platforms, but replication of results has been absent in all but a few cases, and the range of control samples has been limited (Turner Martin R & Benatar Michael, 2015).

The identification of reliable biomarkers of disease could be helpful in clinical practice due to the clinical and genetic heterogeneity of ALS. Biomarkers are critically important to studying pre-symptomatic ALS and essential to efforts to intervene therapeutically before clinically manifest disease emerges (<u>Benatar Michael</u> et al., <u>2022</u>). There is an urgent need to identify biomarkers to detect and study disease progression in ALS <u>(Jeffrey Jeremy</u> et al., <u>2018</u>).

Fluid-based markers from cerebrospinal fluid, blood, and urine are emerging as useful diagnostic, pharmacodynamic, and predictive biomarkers <u>(McMackin Roisin</u> et al., <u>2023</u>). The pursuit of new biomarkers in ALS constitutes a complex process with the potential to impact diagnostics, enhance comprehension of pathogenesis, and contribute to the development of new therapeutic strategies <u>(Klíčová Kateřina</u> et al., <u>2024</u>). Identifying specific biomarkers that aid in establishing disease prognosis, particularly in terms of predicting disease progression, will help our understanding of ALS pathophysiology and could be used to monitor a patient’s response to drugs and therapeutic agents <u>(Waller Rachel</u> et al., <u>2023</u>).

The identification of quantitative and reproducible markers of disease stratification in ALS is fundamental for study design definition and inclusion of homogenous patient cohorts into clinical trials <u>(Spinelli Edoardo G</u> et al., <u>2024</u>). The development of brain imaging biomarkers is essential to advance in the diagnosis, stratification, and monitoring of ALS, both in clinical practice and clinical trials <u>(Mazón Miguel</u> et al., <u>2018</u>). Identifying specific biomarkers that aid in establishing disease prognosis, particularly in terms of predicting disease progression, will help our understanding of ALS pathophysiology and could be used to monitor a patient’s response to drugs and therapeutic agents (<u>Waller Rachel</u> et al., <u>2023</u>). Ultimately, the identification of biomarkers for ALS is crucial for early diagnosis, monitoring disease progression, and evaluating therapeutic responses. Despite significant research efforts, validated biomarkers for clinical use are still lacking, highlighting the need for continued investigation in this area.

Identification of gene variants associated with ALS has informed concepts of the pathogenesis of ALS, aided the identification of therapeutic targets, facilitated research to develop new ALS biomarkers, and supported the establishment of clinical diagnostic tests for ALS-linked genes <u>(Boylan Kevin, 2015</u>). Recent studies have highlighted that there is significant heterogeneity with regard to anatomical and temporal disease progression. Importantly, more recent advances in genetics have revealed new causative genes to the disease. New efforts have focused on the development of biomarkers that could aid in diagnosis, prognosis, and serve as pharmacodynamics markers <u>(Ilieva Hristelina</u> & <u>Maragakis</u> <u>Nicholas J, 2017</u>). Biomarkers that have been widely explored are indispensable for the diagnosis, treatment, and prevention of ALS. The development of new genes and targets is an urgent task in this field (<u>Yang Xiaoming</u> et al., <u>2021</u>). Genetic forms of ALS offer a unique opportunity for therapeutic development, as genetic associations may reveal potential insights into disease etiology. Genetic ALS may also be amenable to investigating earlier intervention given the possibility of identifying clinically presymptomatic, at-risk individuals with causative genetic variants <u>(Benatar Michael</u> et al., <u>2022</u>). Identifying biomarkers that detect these processes during the presymptomatic stage could provide a means to monitor impending disease onset in individuals with genetic mutations associated with familial ALS (fALS) (<u>Irwin</u> <u>Katherine E</u> et al., <u>2024</u>). Overall, our study provides potential biomarkers of ALS for disease diagnosis and therapeutic monitoring <u>(Xie</u> et al., <u>2021</u>). The identification of reliable biomarkers of disease could be helpful in clinical practice <u>(Riolo Giulia</u> et al., <u>2022</u>). Thus, the identification of biomarkers specific for ALS could be of fundamental importance in clinical practice. An ideal biomarker should display high specificity and sensitivity for discriminating ALS from control subjects and from ALS-mimics and other neurological diseases, and should then monitor disease progression within individual patients (<u>Ricci Claudia</u> et al., <u>2018</u>). Therefore, establishing biomarkers for ALS patients is vital for rapid and accurate diagnosis.

Several biomarkers including genetic, biochemical, neurophysiological, cerebrospinal fluid (CSF) and imaging data have been reported for the diagnosis and evaluating progression of ALS, however none of them have been studied in the preclinical stage of ALS and superiority of different biomarkers have not been evaluated until now <u>(Aydemir Duygu &</u> <u>Ulusu Nuriye Nuray, 2020</u>). Uncovering the identity of the genetic factors in ALS will not only improve the accuracy of ALS diagnosis, but may also provide new approaches for preventing and treating the disease (<u>Yamashita & Ando, 2015</u>). The search for protein biomarkers in ALS has been a significant focus in recent research. Fifteen proteins, including APOB, APP, CAMK2A, CHI3L1, CHIT1, CLSTN3, ERAP2, FSTL4, GPNMB, JCHAIN, L1CAM, NPTX2, SERPINA1, SERPINA3, and UCHL1, have shown significant differences between ALS patients and control subjects, providing a foundation for developing new biomarkers for ALS <u>(Oh Sungtaek</u> et al., <u>2023</u>). Additionally, neurofilaments are the most used markers in ALS clinical studies, and other neuroinflammatory-related proteins, p75ECD, p-Tau/t-Tau, and UCHL1, have been tested in multicentered studies and across different laboratories <u>(Donini</u> <u>Luisa</u> et al., <u>2023</u>). Proteins involved in angiogenesis, proteinopathy, and neuroinflammation have shown significantly reduced expression in ALS patients compared to controls, with positive correlations observed among these proteins <u>(Modgil Shweta</u> et al., <u>2020</u>). The identification of reliable protein biomarkers is crucial for early diagnosis, tracking disease progression, and testing target engagement of promising therapeutics (<u>Wilkins Heather M.</u> et al., <u>2021</u>).

Metabolic dysfunction has been suggested to be involved in the pathophysiology of ALS. This study aimed to investigate the potential role of metabolic biomarkers in the progression of ALS and understand the possible metabolic mechanisms. Fifty-two patients with ALS and 24 normal controls were included, and blood samples were collected for analysis of metabolic biomarkers <u>(Li Jin-Yue</u> et al., <u>2022</u>). In this review, we discuss current and potential metabolism biomarkers in the context of ALS. Of those for which data does exist, there is limited insight provided by individual markers, with specificity for disease, and lack of reproducibility and efficacy in informing prognosis being the largest drawbacks <u>(Kirk</u> <u>Siobhan E</u> et al., <u>2019</u>). Metabolomics contribute to a better understanding of ALS pathophysiology but, to date, no biomarker has been validated for diagnosis, principally due to the heterogeneity of the disease and the absence of applied standardized methodology for biomarker discovery. A consensus on best metabolomics methodology as well as systematic independent validation will be an important accomplishment on the path to identifying the long-awaited biomarkers for ALS and to improve clinical trial designs <u>(Blasco H</u> et al., <u>2016</u>). Therefore, the detection of blood biomarkers to be used as screening tools for disease onset and progression has been an expanding research area with few advances in the development of drugs for the treatment of ALS. In this review, we will address the available data analyzing the relationship of lipid metabolism and lipid derived-products with ALS <u>(Trostchansky</u> <u>Andres, 2019</u>). Additionally, we identified novel metabolites to be employed as biomarkers for diagnosis and prognosis of ALS patients (<u>Lanznaster Débora</u> et al., <u>2022</u>). Ultimately, our findings indicate that a panel of metabolites were correlated with disease severity of ALS, which could be potential biomarkers for monitoring ALS progression and therapeutic effects (<u>Chang Kuo-Hsuan</u> et al., <u>2021</u>).

ALS extends beyond the boundaries of the central nervous system, with metabolic alterations being observed at the systemic and cellular level. While the number of studies that assess the role and impact of metabolic perturbations in ALS is rapidly increasing, the use of metabolism biomarkers in ALS remains largely underinvestigated (Kirk Siobhan E et al., 2019). Glucose was increased in CSF of ALS patients and α-hydroxybutyrate was increased in CSF and plasma of ALS patients compared to matched controls. Leucine, isoleucine and ketoleucine were increased in CSF of both ALS and PD. Together, these studies, in conjunction with earlier studies, suggest alterations in energy utilization pathways and have identified and further validated perturbed metabolites to be used in panels of biomarkers for the diagnosis of ALS and PD (Wuolikainen Anna et al., 2016). Our work is helpful for better understanding of metabolic dysfunctions in ALS and provides novel targets for the therapeutic intervention in the future (Li Chunyu et al., 2019). Thus protein, lipid and carbohydrate metabolisms are.

#### Diagnostic Biomarkers in Amyotrophic Lateral Sclerosis

The identification of biomarkers specific for ALS is crucial for clinical practice. An ideal biomarker should display high specificity and sensitivity for discriminating ALS from control subjects and from ALS-mimics and other neurological diseases, and should then monitor disease progression within individual patients. Establishing biomarkers for ALS patients is vital for rapid and accurate diagnosis. Several biomarkers including genetic, biochemical, neurophysiological, cerebrospinal fluid (CSF) and imaging data have been reported for the diagnosis and evaluating progression of ALS <u>(Aydemir Duygu & Ulusu Nuriye Nuray, 2020</u>).

The identification of effective and objective biomarkers may be a feasible method for the early and accurate diagnosis of ALS. Reliable, accessible biomarkers are needed to aid early ALS diagnosis, differentiate from ALS-mimicking diseases, predict survival, and monitor disease progression and treatment response. The identification of biomarkers specific for ALS could be of fundamental importance in clinical practice. An ideal biomarker should exhibit high specificity and sensitivity for distinguishing ALS from control populations and monitor disease progression within individual patients. The development and inclusion of different types of biomarkers in diagnosis and clinical trials can assist in determining target engagement of a drug, in distinguishing between ALS and other diseases, and in predicting disease progression rate, drug responsiveness, or an adverse event. The past decade has seen a dramatic increase in the discovery of candidate biomarkers for ALS. These biomarkers typically can either differentiate ALS from control subjects or predict disease course (slow versus fast progression).

Therefore, new methods are required for the earlier diagnosis of ALS. Screening for pathogenic variants in known ALS-associated genes is already exploited as a diagnostic tool in ALS but cannot be applied for population-based screening. New circulating biomarkers (proteins or small molecules) are needed for initial screening, whereas specific diagnostic methods can be applied to confirm the presence of pathogenic variants in the selected population subgroup (<u>Pampalakis</u> et al., <u>2019</u>).

The ALS pathogenesis related to the diagnostic biomarkers might lessen the diagnostic reliance on the clinical manifestations. Among them, the cortical altered signatures of ALS patients derived from both structural and functional magnetic resonance imaging and the emerging proteomic biomarkers of neuronal loss and glial activation in the cerebrospinal fluid as well as the potential biomarkers in blood, serum, urine, and saliva are leading a new phase of biomarkers. The identification of a panel of biomarkers that accurately reflect features of pathology is a priority, not only for diagnostic purposes but also for prognostic or predictive applications.

Significant progress has been made using cerebrospinal fluid, serum, and plasma in the search for ALS biomarkers, with urine and saliva biomarkers still in earlier stages of development. The development and inclusion of different types of biomarkers in diagnosis and clinical trials can assist in determining target engagement of a drug, distinguishing between ALS and other diseases, and predicting disease progression rate, drug responsiveness, or an adverse event.

Prognostic biomarkers that predict disease severity and progression would be informative for patients with ALS, aiding in care plan development and helping direct patient expectations (Irwin Katherine E et al., 2024). The identification of a panel of biomarkers that accurately reflect features of pathology is a priority, not only for diagnostic purposes but also for prognostic or predictive applications (Kadena & Vlamos, 2021).

The identification of more reliable diagnostic or prognostic biomarkers in age-related neurodegenerative diseases, such as ALS, is urgently needed. The objective in this study was to identify more reliable prognostic biomarkers of ALS mirroring neurodegeneration that could be of help in clinical trials (Calvo Ana C et al., 2019).

Incorporation of biomarkers into the ALS drug development pipeline and the use of biologic and/or imaging biomarkers in early- and late-stage ALS clinical trials have been absent and only recently pursued in early-phase clinical trials. Further clinical research studies are needed to validate biomarkers for disease progression and develop biomarkers that can help determine that a drug has reached its target within the central nervous system (Bakkar Nadine et al., 2015).

The ALS pathogenesis related to the diagnostic biomarkers might lessen the diagnostic reliance on the clinical manifestations. Among them, the cortical altered signatures of ALS patients derived from both structural and functional magnetic resonance imaging and the emerging proteomic biomarkers of neuronal loss and glial activation in the cerebrospinal fluid as well as the potential biomarkers in blood, serum, urine, and saliva are leading a new phase of biomarkers.

Thus, reliable biomarkers of disease activity are urgently needed to stratify patients into homogenous groups with aligned disease trajectories to allow a more effective design of clinical trials. In this review, the most promising candidate biomarkers in the cerebrospinal fluid (CSF) of patients with ALS will be summarised. Correlations between biomarker levels and clinical outcome parameters are discussed, while highlighting potential pitfalls and intercorrelations of these clinical parameters (Dreger Marie et al., 2022).

The identification of biomarkers specific for ALS is crucial for distinguishing ALS from control subjects and other neurological diseases, as well as monitoring disease progression within individual patients (Ricci Claudia et al., 2018). An ideal biomarker should exhibit high specificity and sensitivity for distinguishing ALS from control populations and monitor disease progression within individual patients (Vu Lucas T & Bowser Robert, 2017).

The development and inclusion of different types of biomarkers in diagnosis and clinical trials can assist in distinguishing between ALS and other diseases <u>(Staats Kim A</u> et al., <u>2022</u>). Reliable, accessible biomarkers are needed to aid early ALS diagnosis, differentiate from ALS-mimicking diseases, predict survival, and monitor disease progression and treatment response (<u>Lynch Karen, 2023</u>).

The current diagnosis of ALS mainly depends on clinical manifestations, which contributes to diagnostic delay and difficulty in making an accurate diagnosis at the early stage of ALS (Xu Zheqi & Xu Renshi, 2024). Misdiagnosing ALS can have devastating consequences, including unnecessary emotional burden, delayed and/or inappropriate treatment, and undue financial burden (Lynch Karen, 2023). The identification of effective and objective biomarkers may be a feasible method for the early and accurate diagnosis of ALS (Xu & Yuan, 2021).

#### Prognostic Biomarkers in Amyotrophic Lateral Sclerosis

The identification of biomarkers specific to ALS is also crucial for distinguishing ALS from control subjects and other neurological diseases, as well as monitoring disease progression within individual patients. Prognostic biomarkers that predict disease severity and progression would be informative for patients with ALS, aiding in care plan development and helping direct patient expectations. The development of novel biomarkers for ALS holds the potential to enhance both the diagnosis and comprehension of the often unpredictable disease progression (Klíčová Kateřina et al., 2024).

Several biomarkers, including genetic, biochemical, neurophysiological, cerebrospinal fluid (CSF), and imaging data, have been reported for the diagnosis and evaluation of ALS progression <u>(Aydemir Duygu & Ulusu Nuriye Nuray, 2020</u>). Reliable, accessible biomarkers are needed to aid early ALS diagnosis, differentiate from ALS-mimicking diseases, predict survival, and monitor disease progression and treatment response. The identification of a panel of biomarkers that accurately reflect features of pathology is a priority, not only for diagnostic purposes but also for prognostic or predictive applications (<u>Kadena & Vlamos,</u> <u>2021</u>).

Prognostic biomarkers that predict disease severity and progression would be informative for ALS patients. From a clinical perspective, reliable prognostic biomarkers would aid in care plan development and help direct patient expectations (Irwin Katherine E et al., 2024). Because ALS is increasingly perceived as a multi-system disease, the identification of a panel of biomarkers that accurately reflect features of pathology is a priority, not only for diagnostic purposes but also for prognostic or predictive applications (Vijayakumar Udaya Geetha et al., 2019). The identification of more reliable diagnostic or prognostic biomarkers in age-related neurodegenerative diseases, such as ALS, is urgently needed. The objective in this study was to identify more reliable prognostic biomarkers of ALS mirroring neurodegeneration that could be of help in clinical trials (Calvo Ana C et al., 2019).

Personalized survival prediction models have been proposed to offer a more detailed prognosis for ALS patients (Vidovic Maximilian et al., 2023). The rate of disease progression, measured by the decline of ALS Functional Rating Scale-Revised (ALSFRS-R) from symptom onset to diagnosis (ΔFS), is a well-established prognostic biomarker for predicting survival (Alves Inês et al., 2025). Multiple Cox proportional hazards modelling identified ΔFS as a highly significant independent predictor of survival in ALS (Labra Julie et al., 2016).

Adding immune markers to prediction models dramatically increased short-term prediction compared with routine clinical prognostic variables alone, and the addition of NK-cell markers further improved the prediction accuracy in female participants (Murdock Benjamin J et al., 2024). Specific immune profiles likely contribute to ALS progression in an age and sex-dependent manner, and peripheral immune markers enhance the prediction of short-term clinical outcomes (Murdock Benjamin J et al., 2024).

The ALS-Survival Score (ALS-SS) was developed by assigning scores to various factors, allowing the identification of groups with different survival curves (Kjældgaard Anne-Lene et al., 2021). The ALS-MITOS system can reliably predict the course of ALS up to 18 months and can be considered a novel and valid outcome measure in randomized controlled trials (Tramacere Irene et al., 2015).

Biomarkers have already been used as target markers and outcome parameters for novel treatment approaches in ALS. Several novel biomarkers have shown encouraging results but should be discussed in the context of their early stage of assay and clinical establishment (<u>Witzel Simon</u> et al., <u>2022</u>). It is crucial to further develop and validate ALS biomarkers and incorporate these biomarkers into the ALS drug development process. The development and inclusion of different types of biomarkers in diagnosis and clinical trials can assist in determining target engagement of a drug, in distinguishing between ALS and other diseases, and in predicting disease progression rate, drug responsiveness, or an adverse event.

Further progress remains to be made to find treatment-specific target engagement biomarkers along with readouts of treatment response that can be reliably applied to all emerging therapies and clinical studies (Malaspina Andrea, 2024). Additional progress is still needed for biomarker development and validation to confirm target engagement in ALS treatment trials. Selection of an appropriate biofluid biomarker depends on the treatment mechanism of interest, and biomarker studies should be incorporated into early phase trials (Fournier Christina N, 2022). Inflammatory molecules and proteins may be used as independent predictors of patient survival and might be used in patient stratification and in evaluating the therapeutic response in clinical trials (Jiang Zongzhi et al., 2022).

The identification of effective and objective biomarkers may be a feasible method for the early and accurate diagnosis of ALS. Reliable, accessible biomarkers are needed to aid early ALS diagnosis, differentiate from ALS-mimicking diseases, predict survival, and monitor disease progression and treatment response. The identification of biomarkers specific for ALS could be of fundamental importance in clinical practice. An ideal biomarker should exhibit high specificity and sensitivity for distinguishing ALS from control populations and monitor disease progression within individual patients.

The development and inclusion of different types of biomarkers in diagnosis and clinical trials can assist in determining target engagement of a drug, in distinguishing between ALS and other diseases, and in predicting disease progression rate, drug responsiveness, or an adverse event. The past decade has seen a dramatic increase in the discovery of candidate biomarkers for ALS. These biomarkers typically can either differentiate ALS from control subjects or predict disease course (slow versus fast progression) (<u>Bakkar Nadine</u> et al., <u>2015</u>).

Therefore, new methods are required for the earlier diagnosis of ALS. Screening for pathogenic variants in known ALS-associated genes is already exploited as a diagnostic tool in ALS but cannot be applied for population-based screening. New circulating biomarkers (proteins or small molecules) are needed for initial screening, whereas specific diagnostic methods can be applied to confirm the presence of pathogenic variants in the selected population subgroup <u>(Pampalakis</u> et al., <u>2019</u>). The ALS pathogenesis related to the diagnostic biomarkers might lessen the diagnostic reliance on the clinical manifestations. Among them, the cortical altered signatures of ALS patients derived from both structural and functional magnetic resonance imaging and the emerging proteomic biomarkers of neuronal loss and glial activation in the cerebrospinal fluid as well as the potential biomarkers in blood, serum, urine, and saliva are leading a new phase of biomarkers.

Thus, reliable biomarkers of disease activity are urgently needed to stratify patients into homogenous groups with aligned disease trajectories to allow a more effective design of clinical trials. In this review, the most promising candidate biomarkers in the cerebrospinal fluid (CSF) of patients with ALS will be summarized. Correlations between biomarker levels and clinical outcome parameters are discussed, while highlighting potential pitfalls and intercorrelations of these clinical parameters (Dreger Marie et al., 2022). The identification of biomarkers specific for ALS is crucial for distinguishing ALS from control subjects and other neurological diseases, as well as monitoring disease progression within individual patients (Ricci Claudia et al., 2018). An ideal biomarker should exhibit high specificity and sensitivity for distinguishing ALS from control populations and monitor disease progression within individual patients (Vu Lucas T & Bowser Robert, 2017).

The development and inclusion of different types of biomarkers in diagnosis and clinical trials can assist in distinguishing between ALS and other diseases <u>(Staats Kim A</u> et al., <u>2022</u>). Reliable, accessible biomarkers are needed to aid early ALS diagnosis, differentiate from ALS-mimicking diseases, predict survival, and monitor disease progression and treatment response (<u>Lynch Karen, 2023</u>).

The current diagnosis of ALS mainly depends on clinical manifestations, which contributes to diagnostic delay and difficulty in making an accurate diagnosis at the early stage of ALS (Xu Zheqi & Xu Renshi, 2024). Misdiagnosing ALS can have devastating consequences, including unnecessary emotional burden, delayed and/or inappropriate treatment, and undue financial burden (Lynch Karen, 2023).

#### Conclusion

The identification and validation of biomarkers in ALS are crucial for early diagnosis, monitoring disease progression, and evaluating therapeutic responses. Biomarkers such as neurofilaments, metabolic markers, and genetic variants have shown promise in distinguishing ALS from other neurological conditions and predicting disease trajectory. However, the current practical use of these biomarkers is limited by the lack of standardized methodologies and validation across diverse patient populations. Additionally, the heterogeneity of ALS presents challenges in developing universally applicable biomarkers. Future studies should focus on large-scale, longitudinal research to validate existing biomarkers and discover new ones. Integrating multi-omics approaches, including genomics, proteomics, and metabolomics, could provide a more comprehensive understanding of ALS pathophysiology. Collaborative efforts across research institutions and the inclusion of diverse patient cohorts will be essential in advancing the field. Ultimately, the development of reliable and accessible biomarkers will enhance clinical trial design, improve patient stratification, and facilitate the discovery of effective treatments for ALS.

#### References

Alves Inês, Gromicho Marta, Oliveira Santos Miguel, Pinto Susana & de Carvalho Mamede Assessing disease progression in ALS: prognostic subgroups and outliers. Amyotrophic lateral sclerosis & frontotemporal degeneration (2025).

Aydemir Duygu & Ulusu Nuriye Nuray Importance of the serum biochemical parameters as potential biomarkers for rapid diagnosis and evaluating preclinical stage of ALS. Medical hypotheses (2020).

Bakkar Nadine, Boehringer Ashley & Bowser Robert Use of biomarkers in ALS drug development and clinical trials. Brain research (2015).

Barritt Andrew W, Gabel Matt C, Cercignani Mara & Leigh P Nigel Emerging Magnetic Resonance Imaging Techniques and Analysis Methods in Amyotrophic Lateral Sclerosis. Frontiers in neurology (2018).

Benatar Michael et al. ALS biomarkers for therapy development: State of the field and future directions. Muscle & nerve (2016).

Benatar Michael et al. ALS biomarkers for therapy development: State of the field and future directions. Muscle & nerve (2016).

Benatar Michael et al. Design of a Randomized, Placebo-Controlled, Phase 3 Trial of Tofersen Initiated in Clinically Presymptomatic SOD1 Variant Carriers: the ATLAS Study. Neurotherapeutics : the journal of the American Society for Experimental NeuroTherapeutics (2022). Benatar Michael et al. Preventing amyotrophic lateral sclerosis: insights from pre-symptomatic neurodegenerative diseases. Brain : a journal of neurology (2022).

Bhadra Pratiti et al. Quantitative Proteomics and Differential Protein Abundance Analysis after Depletion of Putative mRNA Receptors in the ER Membrane of Human Cells Identifies Novel Aspects of mRNA Targeting to the ER. Molecules (Basel, Switzerland) (2021).

Blasco H et al. Metabolomics in amyotrophic lateral sclerosis: how far can it take us? European journal of neurology (2016).

Boylan Kevin Familial Amyotrophic Lateral Sclerosis. Neurologic clinics (2015).

Calvo Ana C et al. Collagen XIX Alpha 1 Improves Prognosis in Amyotrophic Lateral Sclerosis. Aging and disease (2019).

Chang Kuo-Hsuan et al. Altered Metabolic Profiles of the Plasma of Patients with Amyotrophic Lateral Sclerosis. Biomedicines (2021).

Costa Júlia & de Carvalho Mamede Emerging molecular biomarker targets for amyotrophic lateral sclerosis. Clinica chimica acta; international journal of clinical chemistry (2016).

Donini Luisa, Tanel Raffaella, Zuccarino Riccardo & Basso Manuela Protein biomarkers for the diagnosis and prognosis of Amyotrophic Lateral Sclerosis. Neuroscience research (2023).

Dreger Marie, Steinbach Robert, Otto Markus, Turner Martin R & Grosskreutz Julian Cerebrospinal fluid biomarkers of disease activity and progression in amyotrophic lateral sclerosis. Journal of neurology, neurosurgery, and psychiatry (2022).

Falzone Yuri Matteo et al. Integrated evaluation of a panel of neurochemical biomarkers to optimize diagnosis and prognosis in amyotrophic lateral sclerosis. European journal of neurology (2022).

Fournier Christina N Considerations for Amyotrophic Lateral Sclerosis (ALS) Clinical Trial Design. Neurotherapeutics : the journal of the American Society for Experimental NeuroTherapeutics (2022).

Ilieva Hristelina & Maragakis Nicholas J Motoneuron Disease: Clinical. Advances in neurobiology (2017).

Irwin Katherine E, Sheth Udit, Wong Philip C & Gendron Tania F Fluid biomarkers for amyotrophic lateral sclerosis: a review. Molecular neurodegeneration (2024).

Jeffrey Jeremy, D’Cunha Hannah & Suzuki Masatoshi Blood Level of Glial Fibrillary Acidic Protein (GFAP) Does not Correlate With Disease Progression in a Rat Model of Familial ALS (SOD1G93A Transgenic). Frontiers in neurology (2018).

Jiang Zongzhi, Wang Ziyi, Wei Xiaojing & Yu Xue-Fan Inflammatory checkpoints in amyotrophic lateral sclerosis: From biomarkers to therapeutic targets. Frontiers in immunology (2022).

Kadena & Vlamos The Importance of Diagnostic and Prognostic Biomarker Identification and Classification Towards Understanding ALS Pathogenesis. Advances in experimental medicine and biology (2021).

Kirk Siobhan E, Tracey Timothy J, Steyn Frederik J & Ngo Shyuan T Biomarkers of Metabolism in Amyotrophic Lateral Sclerosis. Frontiers in neurology (2019).

Kjældgaard Anne-Lene et al. Prediction of survival in amyotrophic lateral sclerosis: a nationwide, Danish cohort study. BMC neurology (2021).

Klíčová Kateřina et al. Utilizing neurodegenerative markers for the diagnostic evaluation of amyotrophic lateral sclerosis. European journal of medical research (2024).

Labra Julie, Menon Parvathi, Byth Karen, Morrison Shea & Vucic Steve Rate of disease progression: a prognostic biomarker in ALS. Journal of neurology, neurosurgery, and psychiatry (2016).

Lanznaster Débora et al. Metabolic Profile and Pathological Alterations in the Muscle of Patients with Early-Stage Amyotrophic Lateral Sclerosis. Biomedicines (2022).

Li Chunyu et al. Decreased Glycogenolysis by miR-338-3p Promotes Regional Glycogen Accumulation Within the Spinal Cord of Amyotrophic Lateral Sclerosis Mice. Frontiers in molecular neuroscience (2019).

Li Jin-Yue et al. Alterations in metabolic biomarkers and their potential role in amyotrophic lateral sclerosis. Annals of clinical and translational neurology (2022).

Lv Yongting & Li Hongfu Blood diagnostic and prognostic biomarkers in amyotrophic lateral sclerosis. Neural regeneration research (2024).

Lynch Karen Pathogenesis and presentation of ALS: examining reasons for delayed diagnosis and identifying opportunities for improvement. The American journal of managed care (2023).

Malaspina Andrea Use of biomarkers in clinical trials and future developments that will help identify novel biomarkers. International review of neurobiology (2024).

Mazón Miguel, Vázquez Costa Juan Francisco, Ten-Esteve Amadeo & Martí-Bonmatí Luis Imaging Biomarkers for the Diagnosis and Prognosis of Neurodegenerative Diseases. The Example of Amyotrophic Lateral Sclerosis. Frontiers in neuroscience (2018).

McMackin Roisin, Bede Peter, Ingre Caroline, Malaspina Andrea & Hardiman Orla Biomarkers in amyotrophic lateral sclerosis: current status and future prospects. Nature reviews. Neurology (2023).

Modgil Shweta, Khosla Radhika, Tiwari Abha, Sharma Kaushal & Anand Akshay Association of Plasma Biomarkers for Angiogenesis and Proteinopathy in Indian Amyotrophic Lateral Sclerosis Patients. Journal of neurosciences in rural practice (2020).

Murdock Benjamin J et al. Peripheral Immune Profiles Predict ALS Progression in an Age- and Sex-Dependent Manner. Neurology(R) neuroimmunology & neuroinflammation (2024).

Oh Sungtaek, Jang Yura & Na Chan Hyun Discovery of Biomarkers for Amyotrophic Lateral Sclerosis from Human Cerebrospinal Fluid Using Mass-Spectrometry-Based Proteomics. Biomedicines (2023).

Pampalakis, et al. New molecular diagnostic trends and biomarkers for amyotrophic lateral sclerosis. Human mutation (2019).

Ricci Claudia, Marzocchi Carlotta & Battistini Stefania MicroRNAs as Biomarkers in Amyotrophic Lateral Sclerosis. Cells (2018).

Riolo Giulia et al. BDNF and Pro-BDNF in Amyotrophic Lateral Sclerosis: A New Perspective for Biomarkers of Neurodegeneration. Brain sciences (2022).

Sanchez-Tejerina Daniel et al. Biofluid Biomarkers in the Prognosis of Amyotrophic Lateral Sclerosis: Recent Developments and Therapeutic Applications. Cells (2023).

Spinelli Edoardo G et al. Structural and Functional Brain Network Connectivity at Different King’s Stages in Patients With Amyotrophic Lateral Sclerosis. Neurology (2024).

Staats Kim A, Borchelt David R, Tansey Malú Gámez & Wymer James Blood-based biomarkers of inflammation in amyotrophic lateral sclerosis. Molecular neurodegeneration (2022).

Sturmey Ellie & Malaspina Andrea Blood biomarkers in ALS: challenges, applications and novel frontiers. Acta neurologica Scandinavica (2022).

Takahashi Ikuko et al. Identification of plasma microRNAs as a biomarker of sporadic Amyotrophic Lateral Sclerosis. Molecular brain (2015).

Tramacere Irene et al. The MITOS system predicts long-term survival in amyotrophic lateral sclerosis. Journal of neurology, neurosurgery, and psychiatry (2015).

Trostchansky Andres Overview of Lipid Biomarkers in Amyotrophic Lateral Sclerosis (ALS). Advances in experimental medicine and biology (2019).

Turner Martin R & Benatar Michael Ensuring continued progress in biomarkers for amyotrophic lateral sclerosis. Muscle & nerve (2015).

Vidovic Maximilian et al. Current State and Future Directions in the Diagnosis of Amyotrophic Lateral Sclerosis. Cells (2023).

Vijayakumar Udaya Geetha et al. A Systematic Review of Suggested Molecular Strata, Biomarkers and Their Tissue Sources in ALS. Frontiers in neurology (2019).

Vu Lucas T & Bowser Robert Fluid-Based Biomarkers for Amyotrophic Lateral Sclerosis. Neurotherapeutics : the journal of the American Society for Experimental NeuroTherapeutics (2017).

Waller Rachel et al. Establishing mRNA and microRNA interactions driving disease heterogeneity in amyotrophic lateral sclerosis patient survival. Brain communications (2023).

Wilkins Heather M., Dimachkie Mazen M. & Agbas Abdulbaki Blood-based Biomarkers for Amyotrophic Lateral Sclerosis Amyotrophic Lateral Sclerosis (2021).

Witzel Simon, Mayer Kristina & Oeckl Patrick Biomarkers for amyotrophic lateral sclerosis. Current opinion in neurology (2022).

Wuolikainen Anna et al. Multi-platform mass spectrometry analysis of the CSF and plasma metabolomes of rigorously matched amyotrophic lateral sclerosis, Parkinson’s disease and control subjects. Molecular bioSystems (2016).

Xie, Luo, He & Tang Novel Insight Into the Role of Immune Dysregulation in Amyotrophic Lateral Sclerosis Based on Bioinformatic Analysis. Frontiers in neuroscience (2021).

Xu & Yuan Considerations on the concept, definition, and diagnosis of amyotrophic lateral sclerosis. Neural regeneration research (2021).

Xu Zheqi & Xu Renshi Current potential diagnostic biomarkers of amyotrophic lateral sclerosis. Reviews in the neurosciences (2024).

Yamashita & Ando Genotype-phenotype relationship in hereditary amyotrophic lateral sclerosis. Translational neurodegeneration (2015).

Yang Xiaoming et al. Amyotrophic Lateral Sclerosis: Molecular Mechanisms, Biomarkers, and Therapeutic Strategies. Antioxidants (Basel, Switzerland) (2021).

## Example 3: Pharmacokinetics analysis

User’s selected topic: Pharmacokinetics analysis of nintedanib

Pharmacokinetics analysis

#### Pharmacokinetics Analysis of Nintedanib

##### Absorption

Nintedanib is a kinase inhibitor and antifibrotic drug approved for treating idiopathic pulmonary fibrosis and various cancers. Its oral bioavailability is approximately 5% (Aiassa Virginia et al., 2023). The drug is administered orally, with rapid absorption and peak plasma concentrations occurring at 1 hour (fasted) and 2 hours (fed) postdose. The terminal phase half-life ranges from 11 to 18 hours, and steady-state concentrations are achieved by 6 days after initiating once-daily dosing (Fediuk Daryl J et al., 2020).

Nintedanib is a synthetic orally active tyrosine kinase inhibitor, primarily targeting receptors of the platelet-derived growth factor, fibroblast growth factor, and vascular endothelial growth factor families, and also affecting other kinases such as Src, Flt-3, LCK, and LYN (Alfei Silvana & Schito Anna Maria, 2020). It is used in the treatment of idiopathic pulmonary fibrosis, chronic fibrosing interstitial lung diseases, and lung cancer (Alfei Silvana & Schito Anna Maria, 2020).

Nintedanib can be administered orally regardless of food, and no clinically relevant drug-drug interactions have been reported (Fogli Stefano et al., 2021). However, its use is associated with a higher risk of adverse events, particularly diarrhea, nausea, vomiting, and weight loss, but it is also associated with a lower risk of cough and dyspnea in patients with idiopathic pulmonary fibrosis and fibrotic interstitial lung diseases (Chen Chao-Hsien et al., 2021).

In clinical trials and real-world studies, dose reductions, treatment interruptions, and the use of anti-diarrheal medications were frequently employed to manage adverse events. Effective management of these adverse events is crucial to minimize their impact, especially in elderly patients or those with advanced disease, who may have a greater rate of treatment discontinuation (Podolanczuk Anna J & Cottin Vincent, 2023).

The absolute bioavailability of 100 mg BIBF 1120 (nintedanib) as a soft gelatin capsule was investigated in a trial (Wind Sven et al., 2019). In the absolute bioavailability trial of healthy volunteers, nintedanib showed a high total clearance (geometric mean 1390mL/min) and a high volume of distribution at steady state (Vss = 1050 L). Urinary excretion of IV nintedanib was about 1% of dose; renal clearance was about 20 mL/min and therefore negligible. There was no deviation from dose proportionality after IV administration in the dose range tested. Absolute bioavailability of oral nintedanib (100 mg capsule) relative to IV dosing (4-hour infusion, 6 mg) was slightly below 5%. Nintedanib was quickly absorbed after oral administration. It underwent rapid and extensive first-pass metabolism and followed at least biphasic disposition kinetics (Dallinger Claudia et al., 2106).

Nintedanib absorption is influenced by its interaction with P-glycoprotein (P-gp). The nintedanib-loaded SEB-β-CD complex significantly decreased the efflux of nintedanib by inhibiting P-gp activity (Lu Rong et al., 2022).

##### Distribution

Nintedanib, a kinase inhibitor and antifibrotic drug, has been extensively studied for its pharmacokinetic properties. One important aspect of its pharmacokinetics is its interaction with plasma proteins. The extent of plasma protein binding significantly influences the drug’s distribution, efficacy, and elimination. Nintedanib exhibits a high degree of plasma protein binding, primarily to albumin and alpha-1-acid glycoprotein. This high binding affinity affects the free concentration of the drug available for therapeutic action and its subsequent distribution to tissues.

The distribution of nintedanib in various tissues has been investigated to understand its therapeutic potential and side effects. A study using UPLC-MS/MS for simultaneous determination of nintedanib and its metabolite BIBF 1202 in different tissues of mice provided valuable insights into its tissue distribution. The study revealed that nintedanib is widely distributed across various tissues, with significant concentrations found in the liver, kidneys, and lungs <u>(Xu, Xiao-wei</u> et al., <u>2015</u>). This extensive tissue penetration is consistent with its therapeutic use in treating idiopathic pulmonary fibrosis and certain cancers, where targeting lung tissue and other organs is essential.

Assessing the ability of nintedanib to cross the blood-brain barrier (BBB) is important for understanding its potential central nervous system (CNS) effects and therapeutic applications in CNS-related conditions. However, there is limited information available regarding the permeability of nintedanib across the BBB. The absence of substantial data on this aspect suggests that further research is needed to determine whether nintedanib can effectively penetrate the CNS and what this might mean for its use in treating CNS disorders.

To summarize, nintedanib exhibits high plasma protein binding, which influences its distribution and therapeutic efficacy. Its extensive tissue penetration, particularly in the liver, kidneys, and lungs, aligns with its clinical applications in treating pulmonary and oncological conditions. However, the limited data on its BBB permeability highlights the need for further studies to fully understand its potential CNS effects and applications.

##### Metabolism

Nintedanib undergoes metabolism primarily through hydroxylation, demethylation, glucuronidation, and acetylation reactions (Fazzini Luca et al., 2022). The main metabolic pathway of nintedanib is esterase-mediated hydrolysis followed by glucuronidation. Nintedanib, used to treat idiopathic pulmonary fibrosis and non-small cell lung cancer, is metabolized to an inactive carboxylate derivative, BIBF1202, via hydrolysis and subsequently by glucuronidation to BIBF1202 acyl-glucuronide (BIBF1202-G). BIBF1202-G contains an ester bond that can be hydrolytically cleaved to BIBF1202. Nintedanib hydrolysis was detected in human liver microsomes (HLMs) but not in small intestinal preparations. CES1 was suggested to be responsible for nintedanib hydrolysis based on experiments using recombinant hydrolases and hydrolase inhibitors, as well as proteomic correlation analysis using 25 individual HLM (Nakashima Shimon et al., 2023).

The primary enzymes involved in the metabolism of nintedanib include carboxylesterase 1 (CES1), which plays a crucial role in the hydrolysis of nintedanib to its inactive carboxylate derivative, BIBF1202 (Nakashima Shimon et al., 2023). This hydrolysis process is followed by glucuronidation, a reaction mediated by UDP-glucuronosyltransferases (UGTs), which further converts BIBF1202 to BIBF1202 acyl-glucuronide (Nakashima Shimon et al., 2023). These enzymatic processes are essential for the drug’s metabolism and subsequent elimination from the body.

Nintedanib is metabolized to a pharmacologically inactive carboxylate derivative, BIBF1202, via hydrolysis and subsequently by glucuronidation to BIBF1202 acyl-glucuronide (BIBF1202-G). Since BIBF1202-G contains an ester bond, it can be hydrolytically cleaved to BIBF1202. Nintedanib hydrolysis was detected in human liver microsomes (HLMs) but not in small intestinal preparations. CES1 was suggested to be responsible for nintedanib hydrolysis according to experiments using recombinant hydrolases and hydrolase inhibitors as well as proteomic correlation analysis using 25 individual HLM (Nakashima Shimon et al., 2023).

The metabolic half-life of nintedanib is approximately 9.5 hours, and the average concentration in the steady state is normally reached in one week, with low concentrations remaining stable for a period longer than one year (Serra López-Matencio José M et al., 2021). This duration indicates the time the drug remains active in the body before being metabolized and eliminated. The terminal phase half-life ranges from 11 to 18 hours, and steady-state concentrations are achieved by 6 days after initiating once-daily dosing (Fediuk Daryl J et al., 2020).

Overall, nintedanib undergoes extensive metabolism primarily through esterase-mediated hydrolysis and subsequent glucuronidation. The major enzymes involved in its metabolism include CES1 and UGTs, which convert nintedanib to its inactive metabolites. The metabolic half-life of nintedanib is approximately 9.5 hours, with steady-state concentrations achieved within one week of dosing. These pharmacokinetic properties are essential for understanding the drug’s efficacy and safety profile in clinical use.

##### Excretion

Nintedanib is primarily eliminated through the bile/faecal pathway, accounting for about 93% of the administered dose, with renal excretion contributing to a low percentage of total clearance, approximately 0.65% (Serra López-Matencio José M et al., 2021). This indicates that the drug is predominantly excreted via the hepatic route, with minimal involvement of the renal system. Understanding the primary route of elimination is important for comprehending the drug’s pharmacokinetics and potential interactions with other medications that may affect liver function.

Renal clearance of nintedanib is relatively low, contributing to only a small fraction of the total drug elimination. This low renal excretion suggests that nintedanib’s clearance is not significantly impacted by renal function, making it a suitable option for patients with varying degrees of renal impairment. However, it is essential to monitor renal function in patients receiving nintedanib to ensure that any potential accumulation of the drug or its metabolites does not occur.

Hepatic clearance plays a major role in the elimination of nintedanib. The drug undergoes extensive metabolism in the liver, primarily through esterase-mediated hydrolysis followed by glucuronidation. The main enzymes involved in this process include carboxylesterase 1 (CES1) and UDP-glucuronosyltransferases (UGTs), which convert nintedanib to its inactive metabolites (Nakashima Shimon et al., 2023). This extensive hepatic metabolism highlights the importance of liver function in the pharmacokinetics of nintedanib.

The half-life of nintedanib is approximately 10-15 hours, which indicates the time required for the drug concentration to reduce by half in the body. This half-life is consistent with the drug’s dosing regimen, allowing for steady-state concentrations to be achieved within one week of dosing. The terminal phase half-life ranges from 11 to 18 hours, and steady-state concentrations are typically reached by 6 days after initiating once-daily dosing (Fediuk Daryl J et al., 2020).

Nintedanib has shown efficacy in slowing the ongoing degradation of lung function, reducing hospitalization risk, and delaying exacerbations among patients with idiopathic pulmonary fibrosis (IPF) (Guo Hehua et al., 2023). However, these pharmacological interventions do not present a definitive solution to IPF, positioning lung transplantation as the only potential curative measure in contemporary medical practice. Combining nintedanib with other treatments has demonstrated no safety concerns following 12 weeks of management in patients with primarily mild to moderate IPF (Hadi Dalia D et al., 2023).

In summary, nintedanib undergoes extensive hepatic metabolism and is primarily eliminated through the bile/faecal pathway, with minimal renal excretion. The drug’s half-life of approximately 10-15 hours allows for steady-state concentrations to be achieved within one week of dosing. These pharmacokinetic properties are crucial for understanding the drug’s efficacy and safety profile in clinical use. The potential for drug accumulation with repeated dosing appears to be low, given the drug’s elimination pathways and half-life. This understanding of nintedanib’s pharmacokinetics is essential for optimizing its therapeutic use and managing potential adverse effects.

##### Dosing

Nintedanib, a tyrosine kinase inhibitor, is primarily used in the treatment of idiopathic pulmonary fibrosis (IPF) and certain types of cancer. The recommended dose of nintedanib for IPF, as per clinical guidelines, is 150 mg taken orally twice daily. This dosage has been established based on clinical trials and is aimed at maximizing therapeutic efficacy while minimizing adverse effects (Hartman Stan J F et al., 2020).

The dosing frequency of nintedanib is typically twice daily, which aligns with its pharmacokinetic profile. This regimen helps maintain steady-state plasma concentrations, ensuring continuous inhibition of the target kinases. The twice-daily dosing schedule is supported by pharmacokinetic studies that demonstrate a half-life of approximately 9 to 12 hours for nintedanib, necessitating regular administration to sustain its therapeutic levels (Wind Sven et al., 2019).

Dose adjustments for nintedanib are often required based on patient-specific factors, particularly in response to adverse events. In a real-world retrospective observational study of 224 IPF cases treated with nintedanib, 55.7% of patients experienced adverse events, leading to dose reductions in 28.3% of patients and discontinuation in 13.1% of patients (Chen Chao-Hsien et al., 2021). Common adverse events include gastrointestinal issues such as diarrhea, nausea, and vomiting, which often necessitate dose modifications or temporary treatment interruptions.

In clinical trials, strategies to manage adverse events included dose reductions, treatment interruptions, and the use of supportive medications such as anti-diarrhoeal agents (Podolanczuk Anna J & Cottin Vincent, 2023). These measures are essential for maintaining patient adherence to the treatment regimen and ensuring the continued therapeutic benefit of nintedanib.

Overall, the pharmacokinetic analysis of nintedanib highlights the importance of individualized dosing strategies to optimize treatment outcomes. The recommended dose of 150 mg twice daily is generally effective, but adjustments are often necessary to manage adverse events and accommodate patient-specific needs. This approach ensures that patients receive the maximum benefit from nintedanib while minimizing the risk of treatment-related complications.

##### Mechanism of Action

Nintedanib is a tyrosine kinase inhibitor that targets multiple receptor pathways, including the vascular endothelial growth factor receptor (VEGFR), fibroblast growth factor receptor (FGFR), and platelet-derived growth factor receptor (PDGFR) <u>(Schramm</u> et al., <u>2022</u>; <u>Cantini Luca</u> et al., <u>2020</u>; <u>Solinc Julien</u> et al., <u>2022</u>). It disrupts signaling pathways involved in fibroblast proliferation, migration, and differentiation, as well as the secretion of extracellular matrix (ECM) <u>(Bazdyrev Evgeny</u> et al., <u>2022</u>). Nintedanib also exerts anti-inflammatory activity by inhibiting IL-1β <u>(Bazdyrev Evgeny</u> et al., <u>2022</u>). It has been shown to reduce the decline in forced vital capacity (FVC) in idiopathic pulmonary fibrosis (IPF) patients, suggesting a therapeutic impact on disease progression <u>(Bezerra Frank</u> <u>Silva</u> et al., <u>2023</u>).

Nintedanib is a tyrosine kinase inhibitor that targets multiple receptors involved in fibrotic and inflammatory processes. It has demonstrated efficacy and safety in idiopathic pulmonary fibrosis (IPF), systemic sclerosis-associated interstitial lung disease (SSc-ILD), and other fibrosing interstitial lung diseases (ILDs) with a progressive phenotype <u>(Spagnolo</u> et al., <u>2021</u>). Nintedanib targets fibroblast growth factor receptors (FGFRs), platelet-derived growth factor receptors (PDGFRs), and vascular endothelial growth factor receptors (VEGFRs) (<u>Zhang Hao</u> et al., <u>2022</u>). It is approved by the US Food and Drug Administration (FDA) for slowing the rate of decline in pulmonary function in patients with SSc-ILD <u>(Khanna</u> <u>Dinesh</u> et al., <u>2022</u>). Nintedanib has also been shown to be effective in reducing the decline of forced vital capacity in rheumatoid arthritis-associated interstitial lung disease (RA-ILD) (<u>Akiyama Mitsuhiro & Kaneko Yuko, 2022</u>).

Nintedanib is an oral, small-molecule tyrosine kinase inhibitor approved for the treatment of idiopathic pulmonary fibrosis and patients with advanced non-small cell cancer of adenocarcinoma tumour histology. It competitively binds to the kinase domains of vascular endothelial growth factor (VEGF), platelet-derived growth factor (PDGF), and fibroblast growth factor (FGF) (<u>Wind Sven</u> et al., <u>2019</u>; <u>Wind Sven</u> et al., <u>2019</u>). Nintedanib selectively inhibits the activation and tumour-promoting effects of fibroblasts from lung adenocarcinoma patients (<u>Gabasa M</u> et al., <u>2017</u>).

Studies have shown that nintedanib has time-independent pharmacokinetic characteristics, with maximum plasma concentrations reached approximately 2-4 hours after oral administration (<u>Wind Sven</u> et al., <u>2019</u>). There is no relevant pharmacokinetic drug-drug interaction between nintedanib and pirfenidone <u>(Yvette Lamb, 2021</u>). The relationship between nintedanib exposure, patient characteristics, and clinical efficacy in patients with idiopathic pulmonary fibrosis has been explored, as well as the relationship between nintedanib exposure and adverse events in these patients (<u>Wind Sven</u> et al., <u>2019</u>).

Nintedanib’s pharmacodynamic effects are closely tied to its mechanism of action. By inhibiting key tyrosine kinases, nintedanib interferes with the signaling pathways that promote fibrosis and inflammation. This inhibition results in decreased fibroblast activity, reduced extracellular matrix production, and diminished inflammatory responses. These effects collectively contribute to the slowing of disease progression in conditions like idiopathic pulmonary fibrosis and systemic sclerosis-associated interstitial lung disease.

The therapeutic benefits of nintedanib’s mechanism of action are substantial. By targeting multiple pathways involved in fibrosis and inflammation, nintedanib provides a comprehensive approach to managing fibrotic diseases. Its ability to slow the decline in lung function in patients with idiopathic pulmonary fibrosis and systemic sclerosis-associated interstitial lung disease highlights its potential to alter the course of these debilitating conditions. Its efficacy in reducing forced vital capacity decline in rheumatoid arthritis-associated interstitial lung disease demonstrates its broader applicability in fibrotic lung diseases.

Nintedanib’s pharmacokinetic and pharmacodynamic profiles, along with its therapeutic benefits, make it a valuable treatment option for patients with fibrotic lung diseases. Its ability to target multiple receptor pathways and disrupt key signaling mechanisms involved in fibrosis and inflammation positions it as an essential tool in managing these challenging conditions.

##### Pharmacokinetic Variability

Nintedanib’s pharmacokinetics can be influenced by various factors, including genetic polymorphisms, age, renal and hepatic function, and drug-drug interactions. Understanding these factors is crucial for optimizing its therapeutic use and managing potential adverse effects.

Genetic polymorphisms can significantly impact the metabolism of nintedanib. Variations in genes encoding for enzymes such as carboxylesterase 1 (CES1) and UDP-glucuronosyltransferases (UGTs) may alter the drug’s metabolic rate and efficacy. However, specific studies detailing the influence of genetic polymorphisms on nintedanib metabolism are limited, indicating a need for further research in this area.

Age-related differences in pharmacokinetics are also important to consider. As individuals age, physiological changes can affect drug absorption, distribution, metabolism, and excretion. For nintedanib, age-related pharmacokinetic variations have not been extensively studied, but it is known that elderly patients may experience different drug exposure levels compared to younger individuals. This could be due to age-related decline in liver and kidney function, which are critical for drug metabolism and clearance.

Renal impairment can affect the clearance of nintedanib, although its renal excretion is relatively low. Studies have shown that nintedanib and ripretinib carry the lowest risks of renal impairment among tyrosine kinase inhibitors <u>(Xiong Ying</u> et al., <u>2022</u>). Elevated creatinine and protein levels were the most common nephrotoxic events, whereas hematuria was relatively rare. This suggests that while renal function should be monitored, nintedanib’s clearance is not significantly impacted by renal impairment, making it a suitable option for patients with varying degrees of renal function.

Hepatic impairment plays a more significant role in the pharmacokinetics of nintedanib. The drug undergoes extensive metabolism in the liver, primarily through esterase-mediated hydrolysis followed by glucuronidation. The main enzymes involved in this process include CES1 and UGTs, which facilitate the conversion of nintedanib to its inactive metabolites <u>(Nakashima Shimon</u> et al., <u>2023</u>). Patients with hepatic impairment may experience altered drug metabolism and clearance, necessitating dose adjustments and careful monitoring.

Drug-drug interactions are another critical factor affecting nintedanib’s pharmacokinetics. Administering nintedanib with add-on pirfenidone is supposed to enhance the therapeutic benefit by simultaneously acting on two different pathogenic pathways. However, little information is available about their drug-drug interaction, which is important mainly in polymedicated patients (<u>Serra López-Matencio José M</u> et al., <u>2021</u>). The aim of this review is to describe the current management of progressive fibrosing interstitial lung diseases (PF-ILDs) in general and of IPF in particular, focusing on the pharmacokinetic drug-drug interactions between these two drugs and their relationship with other medications in patients with IPF (<u>Serra López-Matencio José M</u> et al., <u>2021</u>). The discovery of antifibrotic agents has resulted in advances in the therapeutic management of idiopathic pulmonary fibrosis (IPF). Currently, nintedanib and pirfenidone have become the basis of IPF therapy based on the results of large randomized clinical trials showing their safety and efficacy in reducing disease advancement. However, the goal of completely halting disease progress has not been reached yet (<u>Serra López-Matencio José M</u> et al., <u>2021</u>). However, there was a drug-drug interaction between pirfenidone and PBI-4050, while the pharmacokinetic profiles were similar among patients with PBI-4050 alone or in conjunction with nintedanib, encouraging further study of PBI-4050 alone or PBI-4050 plus nintedanib in patients with IPF (<u>Yu</u> et al., <u>2022</u>).

In summary, nintedanib’s pharmacokinetics are influenced by genetic polymorphisms, age, renal and hepatic function, and drug-drug interactions. Genetic variations in enzymes such as CES1 and UGTs may alter the drug’s metabolism, while age-related changes can affect drug exposure levels. Renal impairment has a minimal impact on nintedanib clearance, but hepatic impairment significantly affects its metabolism and clearance. Drug-drug interactions, particularly with pirfenidone, require careful consideration in polymedicated patients. Understanding these factors is essential for optimizing nintedanib’s therapeutic use and managing potential adverse effects.

##### Preclinical Data

Nintedanib has been extensively studied in both in-vitro and in-vivo settings to understand its pharmacokinetic properties and therapeutic potential. In-vitro studies have shown that nintedanib effectively inhibits the growth of tumors formed by FGFR1-amplified NCI-H520 and LK-02 cells <u>(Pekarek Leonel</u> et al., <u>2022</u>). It has also been demonstrated to inhibit angiogenesis and suppress tumor cell proliferation <u>(Yumura Masako</u> et al., <u>2020</u>). Nintedanib also inhibits the phosphorylation of PDGFR and VEGFR, suppresses cell proliferation, and decreases the content of collagen-I and the expression of fibrosis-related genes, thereby protecting against fibrosis in both murine and human PCKS <u>(Qin Rui</u> et al., <u>2022</u>).

Further in-vitro studies have shown that nintedanib inhibits keloid fibroblast function by targeting the MAPK pathway components p38, ERK, and JNK, as well as the phosphorylation of Smad2 and Smad3, in a dose-dependent manner <u>(Xu Wanyu</u> et al., <u>2024</u>). In a cellular model of systemic sclerosis-related interstitial lung disease (SSc-ILD), lung fibroblasts stimulated by PDGF in the presence of nintedanib showed a significant decrease in the production of collagen and fibronectin, and a reduction in the proliferation rate by 1.9-fold within 24 hours (<u>Kafle Sunam</u> et al., <u>2021</u>).

In-vivo studies have also provided valuable insights into the pharmacokinetics and therapeutic efficacy of nintedanib. It has shown positive effects in preventing the decline of lung function in patients with systemic sclerosis-related interstitial lung disease (van den Hoogen Lucas L & van Laar Jacob M, 2020). Nintedanib is approved in Europe for progressive interstitial lung disease in systemic sclerosis (Hellmich Bernhard & Henes Joerg C, 2022). It is a small-molecule oral drug approved by the FDA for its ability to retard the rate of progression of idiopathic pulmonary fibrosis (Yu Zuxiang et al., 2023).

In an animal model of liver fibrosis, nintedanib demonstrated significant therapeutic effects (<u>Wollin Lutz</u> et al., <u>2020</u>). Pharmacokinetic studies in healthy volunteers and patients with advanced cancer have shown that nintedanib has time-independent pharmacokinetic characteristics, with maximum plasma concentrations reached approximately 2-4 hours after oral administration (<u>Wind Sven</u> et al., <u>2019</u>). At a dose of 200 mg twice daily, nintedanib does not exhibit proarrhythmic potential <u>(Wind Sven</u> et al., <u>2019</u>). The metabolism and excretion of nintedanib involve CYP isozymes and glucuronidation (UGT) enzymes (<u>Wind Sven</u> et al., <u>2019</u>).

The geometric mean plasma concentration-time profile of nintedanib after single-dose administration (100 mg) to healthy volunteers shows the pharmacokinetic behavior of the drug (Wind Sven et al., 2019). Consistent pharmacokinetic properties have been observed in both healthy volunteers and patients with advanced cancer (Dallinger Claudia et al.; Wind Sven et al., 2019).

Overall, the preclinical data on nintedanib highlight its potential as a therapeutic agent in various fibrotic and oncologic conditions. The in-vitro and in-vivo studies collectively provide a detailed understanding of its pharmacokinetic properties, mechanisms of action, and therapeutic efficacy.

##### Clinical Data

Nintedanib has shown efficacy in reducing the progression of idiopathic pulmonary fibrosis (IPF) in two phase 3 clinical trials, INPULSIS-1 and INPULSIS-2 (Confalonieri Paola et al., 2022). The INBUILD trial confirmed that antifibrotic treatment with nintedanib is beneficial in other interstitial lung diseases (ILDs) characterized by a progressive phenotype (Albera Carlo et al., 2021).

In the INBUILD trial, nintedanib was shown to prevent worsening of cough in patients with progressive fibrosing interstitial lung diseases, although no significant effect on cough was observed in the INPULSIS trials with IPF patients (Kašiković Lečić Svetlana et al., 2022). Nintedanib has also been tested in combination with other treatments. For instance, the combination of nintedanib with docetaxel provided clinical benefits over docetaxel alone as a second-line treatment for patients with advanced non-small cell lung cancer (NSCLC) of adenocarcinoma histology (Yan Shu et al., 2023). The VARGADO trial reported improved overall response rate (ORR) and median progression-free survival (PFS) in patients with NSCLC receiving second-line nintedanib plus docetaxel after chemoimmunotherapy (Garon Edward B et al., 2023).

In a phase II clinical study, nintedanib showed promising results in recurrent high-grade gliomas, although another phase II trial reported minimal clinical anti-tumor activity despite a good safety profile (Shamshiripour Parisa et al., 2022). Nintedanib has been approved for use in patients with systemic sclerosis-associated interstitial lung disease (SSc-ILD) and progressive fibrosing interstitial lung diseases (PF-ILD) based on the results of the INBUILD trial (Shumar John N et al., 2021).

Nintedanib is a synthetic orally active tyrosine kinase inhibitor that primarily inhibits the receptors of the platelet-derived growth factor, fibroblast growth factor, and vascular endothelial growth factor families. It also affects other kinases, including Src, Flt-3, LCK, and LYN (Grześk Grzegorz et al., 2020). Pharmacokinetic studies have shown that nintedanib is rapidly absorbed after oral administration, with its pharmacokinetics being time-independent and dose-linear (Pan Lei et al., 2023).

Nintedanib has very low oral bioavailability (4.7%), necessitating high doses (100-150 mg) which can cause gastrointestinal irritation and hepatotoxicity <u>(Thakkar Dhruti</u> et al., <u>2024</u>). The pharmacokinetics of nintedanib are consistent between healthy volunteers and patients with IPF, with 95% of the steady-state concentration being reached after approximately five administrations at 18 mg twice a day (<u>Kolb Martin</u> et al., <u>2023</u>).

The co-administration of nintedanib and pirfenidone has shown no relevant effects on pharmacokinetic drug-drug interactions, although clinical trials are still needed to assess whether this combination can improve efficacy (Chen I-Chen et al., 2022). The pharmacokinetic profiles were similar among patients with PBI-4050 alone or in conjunction with nintedanib, encouraging further study of PBI-4050 plus nintedanib in patients with IPF (Yu et al., 2022).

Nintedanib was associated with a higher risk of adverse events, especially diarrhea, nausea, vomiting, and weight loss, but it was also associated with a lower risk of cough and dyspnea in IPF and fibrotic-ILD patients <u>(Chen Chao-Hsien</u> et al., <u>2021</u>). The use of nintedanib was positively associated with diarrhea (OR = 5.96; 95% CI = 4.35-8.16), nausea (OR = 3.00; 95% CI = 1.93-4.66), vomiting (OR = 3.22; 95% CI = 2.17-4.76), and weight loss (OR = 3.38; 95% CI = 1.1.76-6.47). Conversely, patients treated with nintedanib were less likely to have a cough (OR = 0.73; 95% CI = 0.56-0.96) and dyspnea (OR = 0.70; 95% CI = 0.53-0.94) (<u>Chen Chao-Hsien</u> et al., <u>2021</u>).

Caution should be exercised before prematurely endorsing and applying nintedanib therapy. Studies are encouraged to clarify and justify its potential role as switch vs. add-on therapy and to define disease progression more rigorously (<u>Mura Marco, 2021</u>).

##### Summary

Nintedanib, a tyrosine kinase inhibitor, exhibits a complex pharmacokinetic profile that is crucial for its therapeutic efficacy in treating idiopathic pulmonary fibrosis (IPF) and certain cancers. The key pharmacokinetic parameters of nintedanib include its low oral bioavailability of approximately 5%, rapid absorption with peak plasma concentrations occurring within 2-4 hours post-administration, and a half-life ranging from 9.5 to 18 hours. These parameters necessitate a twice-daily dosing regimen to maintain therapeutic levels and ensure continuous inhibition of target kinases.

Absorption and distribution significantly impact the efficacy of nintedanib. Despite its low bioavailability, nintedanib is rapidly absorbed and widely distributed across various tissues, including the liver, kidneys, and lungs. This extensive tissue penetration aligns with its clinical applications in treating pulmonary and oncological conditions. High plasma protein binding, primarily to albumin and alpha-1-acid glycoprotein, influences the drug’s distribution and free concentration available for therapeutic action. The ability of nintedanib to reach and maintain effective concentrations in target tissues is essential for its antifibrotic and antineoplastic effects.

Nintedanib undergoes extensive metabolism, primarily through esterase-mediated hydrolysis followed by glucuronidation. The main enzymes involved in its metabolism include carboxylesterase 1 (CES1) and UDP-glucuronosyltransferases (UGTs). These metabolic pathways convert nintedanib to its inactive metabolites, BIBF1202 and BIBF1202 acyl-glucuronide. Understanding these pathways is crucial for predicting drug interactions and optimizing dosing regimens. While nintedanib has shown minimal clinically relevant drug-drug interactions, its combination with other treatments, such as pirfenidone, requires careful consideration to avoid potential adverse effects.

Elimination of nintedanib is primarily through the bile/faecal pathway, accounting for about 93% of the administered dose, with renal excretion contributing minimally to total clearance. This hepatic route of elimination highlights the importance of liver function in the pharmacokinetics of nintedanib. The low renal excretion suggests that nintedanib’s clearance is not significantly impacted by renal function, making it suitable for patients with varying degrees of renal impairment. However, hepatic impairment can affect the drug’s metabolism and clearance, necessitating dose adjustments and careful monitoring.

The pharmacokinetic properties of nintedanib support the clinical dosing regimen of 150 mg taken orally twice daily. This regimen is designed to maintain steady-state plasma concentrations, ensuring continuous inhibition of target kinases and maximizing therapeutic efficacy. The twice-daily dosing schedule aligns with the drug’s half-life and absorption characteristics, allowing for effective management of IPF and certain cancers. Dose adjustments and treatment interruptions are often employed to manage adverse events, particularly gastrointestinal issues, which are common with nintedanib therapy.

Despite the comprehensive understanding of nintedanib’s pharmacokinetics, several uncertainties remain that warrant further study. The impact of genetic polymorphisms on the metabolism and efficacy of nintedanib is not well understood and requires additional research. Age-related pharmacokinetic variations and the drug’s ability to cross the blood-brain barrier are also areas that need further investigation. Additionally, the long-term effects of nintedanib on liver function and potential drug-drug interactions in polymedicated patients should be explored to optimize its therapeutic use and minimize adverse effects.

Overall, nintedanib’s pharmacokinetic profile, characterized by rapid absorption, extensive tissue distribution, and hepatic metabolism, supports its clinical efficacy in treating IPF and certain cancers. The twice-daily dosing regimen is effective in maintaining therapeutic levels, although dose adjustments are often necessary to manage adverse events. Further research is needed to address uncertainties related to genetic polymorphisms, age-related variations, and long-term safety, which will enhance the understanding and optimization of nintedanib therapy.

##### References

Aiassa Virginia, Garnero Claudia, Zoppi Ariana & Longhi Marcela R Cyclodextrins and Their Derivatives as Drug Stability Modifiers. Pharmaceuticals (Basel, Switzerland) (2023).

Akiyama Mitsuhiro & Kaneko Yuko Pathogenesis, clinical features, and treatment strategy for rheumatoid arthritis-associated interstitial lung disease. Autoimmunity reviews (2022).

Albera Carlo et al. Progressive Fibrosing Interstitial Lung Diseases: A Current Perspective. Biomedicines (2021).

Alfei Silvana & Schito Anna Maria Positively Charged Polymers as Promising Devices against Multidrug Resistant Gram-Negative Bacteria: A Review. Polymers (2020).

Bazdyrev Evgeny et al. The Hidden Pandemic of COVID-19-Induced Organizing Pneumonia. Pharmaceuticals (Basel, Switzerland) (2022).

Bezerra Frank Silva, et al. Oxidative Stress and Inflammation in Acute and Chronic Lung Injuries. Antioxidants (Basel, Switzerland) (2023).

Cantini Luca, Hassan Raffit, Sterman Daniel H & Aerts Joachim G J V Emerging Treatments for Malignant Pleural Mesothelioma: Where Are We Heading? Frontiers in oncology (2020).

Chen Chao-Hsien et al. The safety of nintedanib for the treatment of interstitial lung disease: A systematic review and meta-analysis of randomized controlled trials. PloS one (2021).

Chen I-Chen, et al. Evaluation of Proteasome Inhibitors in the Treatment of Idiopathic Pulmonary Fibrosis. Cells (2022).

Confalonieri Paola et al. Regeneration or Repair? The Role of Alveolar Epithelial Cells in the Pathogenesis of Idiopathic Pulmonary Fibrosis (IPF). Cells (2022).

Dallinger, Claudia et al. “Pharmacokinetic Properties of Nintedanib in Healthy Volunteers and Patients With Advanced Cancer.” Journal of clinical pharmacologyvol. 56,11. (2016).

Fazzini Luca et al. Metabolomic Profiles on Antiblastic Cardiotoxicity: New Perspectives for Early Diagnosis and Cardioprotection. Journal of clinical medicine (2022).

Fediuk Daryl J et al. Overview of the Clinical Pharmacology of Ertugliflozin, a Novel Sodium-Glucose Cotransporter 2 (SGLT2) Inhibitor. Clinical pharmacokinetics (2020).

Fogli Stefano et al. Clinical pharmacology and drug-drug interactions of lenvatinib in thyroid cancer. Critical reviews in oncology/hematology (2021).

Gabasa M, Ikemori R, Hilberg F, Reguart N & Alcaraz J Nintedanib selectively inhibits the activation and tumour-promoting effects of fibroblasts from lung adenocarcinoma patients. British journal of cancer (2017).

Garon Edward B et al. Clinical outcomes of ramucirumab plus docetaxel in the treatment of patients with non-small cell lung cancer after immunotherapy: a systematic literature review. Frontiers in oncology (2023).

Grześk Grzegorz et al. The Interactions of Nintedanib and Oral Anticoagulants-Molecular Mechanisms and Clinical Implications. International journal of molecular sciences (2020).

Guo Hehua et al. Progress in understanding and treating idiopathic pulmonary fibrosis: recent insights and emerging therapies. Frontiers in pharmacology (2023).

Hadi Dalia D et al. Idiopathic pulmonary fibrosis: Addressing the current and future therapeutic advances along with the role of Sotatercept in the management of pulmonary hypertension. Immunity, inflammation and disease (2023).

Hartman Stan J F, et al. Pharmacokinetics and Target Attainment of Antibiotics in Critically Ill Children: A Systematic Review of Current Literature. Clinical pharmacokinetics (2020).

Hellmich Bernhard & Henes Joerg C [Biologics for connective tissue diseases and vasculitides]. Der Internist (2022).

Kafle Sunam, Thapa Magar Manusha, Patel Priyanka, Poudel Arisa & Cancarevic Ivan Systemic Sclerosis Associated Interstitial Lung Disease and Nintedanib: A Rare Disease and a Promising Drug. Cureus (2021).

Kašiković Lečić Svetlana et al. Management of musculoskeletal pain in patients with idiopathic pulmonary fibrosis: a review. Upsala journal of medical sciences (2022).

Khanna Dinesh et al. Systemic Sclerosis-Associated Interstitial Lung Disease: How to Incorporate Two Food and Drug Administration-Approved Therapies in Clinical Practice. Arthritis & rheumatology (Hoboken, N.J.) (2022).

Kolb Martin, Crestani Bruno & Maher Toby M Phosphodiesterase 4B inhibition: a potential novel strategy for treating pulmonary fibrosis. European respiratory review : an official journal of the European Respiratory Society (2023).

Lamb, Yvette N. “Nintedanib: A Review in Fibrotic Interstitial Lung Diseases.” Drugs vol. 81,5 (2021)

Lu Rong, Zhou Yun, Ma Jinqian, Wang Yuchen & Miao Xiaoqing Strategies and Mechanism in Reversing Intestinal Drug Efflux in Oral Drug Delivery. Pharmaceutics (2022).

Mura Marco Use of nintedanib in interstitial lung disease other than idiopathic pulmonary fibrosis: much caution is warranted. Pulmonary pharmacology & therapeutics (2021).

Nakashima, Shimon et al. “Characterization of Enzymes Involved in Nintedanib Metabolism in Humans.” Drug metabolism and disposition: the biological fate of chemicals vol. 51,6 (2023).

Pan Lei et al. Nintedanib in an elderly non-small-cell lung cancer patient with severe steroid-refractory checkpoint inhibitor-related pneumonitis: A case report and literature review. Frontiers in immunology (2023).

Pekarek Leonel et al. Clinical Applications of Classical and Novel Biological Markers of Pancreatic Cancer. Cancers (2022).

Podolanczuk Anna J & Cottin Vincent A Narrative Review of Real-World Data on the Safety of Nintedanib in Patients with Idiopathic Pulmonary Fibrosis. Advances in therapy (2023).

Qin Rui et al. Indole-Based Small Molecules as Potential Therapeutic Agents for the Treatment of Fibrosis. Frontiers in pharmacology (2022).

Schramm, Schaefer & Wygrecka EGFR Signaling in Lung Fibrosis. Cells (2022).

Serra López-Matencio José M, et al. Pharmacological Interactions of Nintedanib and Pirfenidone in Patients with Idiopathic Pulmonary Fibrosis in Times of COVID-19 Pandemic. Pharmaceuticals (Basel, Switzerland) (2021).

Shamshiripour Parisa et al. Next-Generation Anti-Angiogenic Therapies as a Future Prospect for Glioma Immunotherapy; From Bench to Bedside. Frontiers in immunology (2022).

Shumar John N, Chandel Abhimanyu & King Christopher S Antifibrotic Therapies and Progressive Fibrosing Interstitial Lung Disease (PF-ILD): Building on INBUILD. Journal of clinical medicine (2021).

Solinc Julien et al. The Platelet-Derived Growth Factor Pathway in Pulmonary Arterial Hypertension: Still an Interesting Target? Life (Basel, Switzerland) (2022).

Spagnolo, et al. Mechanisms of progressive fibrosis in connective tissue disease (CTD)-associated interstitial lung diseases (ILDs). Annals of the rheumatic diseases (2021).

Thakkar Dhruti, Singh Sanskriti & Wairkar Sarika Advanced Delivery Strategies of Nintedanib for Lung Disorders and Beyond: A Comprehensive Review. AAPS PharmSciTech (2024).

Wind Sven et al. Clinical Pharmacokinetics and Pharmacodynamics of Nintedanib. Clinical pharmacokinetics (2019).

Wollin Lutz, Togbe Dieudonnée & Ryffel Bernhard Effects of Nintedanib in an Animal Model of Liver Fibrosis. BioMed research international (2020).

Xiong Ying, Wang Qinxuan, Liu Yangyi, Wei Jingwen & Chen Xiaolei Renal adverse reactions of tyrosine kinase inhibitors in the treatment of tumours: A Bayesian network meta-analysis. Frontiers in pharmacology (2022).

Xu Wanyu et al. Acne-induced pathological scars: pathophysiology and current treatments. Burns & trauma (2024).

Xu, Xiao-wei et al. “Simultaneous determination of nintedanib and its metabolite BIBF 1202 in different tissues of mice by UPLC-MS/MS and its application in drug tissue distribution study.” Journal of chromatography. B, Analytical technologies in the biomedical and life sciences vol. 1002 (2015).

Yan Shu et al. Efficacy and safety of nintedanib in patients with non-small cell lung cancer, and novel insights in radiation-induced lung toxicity. Frontiers in oncology (2023).

Yu Zuxiang et al. Tissue fibrosis induced by radiotherapy: current understanding of the molecular mechanisms, diagnosis and therapeutic advances. Journal of translational medicine (2023).

Yu et al. Ongoing Clinical Trials in Aging-Related Tissue Fibrosis and New Findings Related to AhR Pathways. Aging and disease (2022).

Yumura Masako, Nagano Tatsuya & Nishimura Yoshihiro Novel Multitarget Therapies for Lung Cancer and Respiratory Disease. Molecules (Basel, Switzerland) (2020).

Zhang Hao, Ren Lu & Shivnaraine Rabindra Vishwadev Targeting GPCRs to treat cardiac fibrosis. Frontiers in cardiovascular medicine (2022).

van den Hoogen Lucas L & van Laar Jacob M Targeted therapies in systemic sclerosis, myositis, antiphospholipid syndrome, and Sjögren’s syndrome. Best practice & research. Clinical rheumatology (2020).

## Example 4: Scientific blog post

User’s selected topic: How COVID-19 changed the future of medicine

Scientific blog post

#### How COVID-19 Revolutionized Medicine: Telehealth, Vaccines, and Beyond

##### Abstract

The COVID-19 pandemic has significantly transformed the field of medicine, driving rapid advancements and necessitating major adaptations across various aspects of healthcare. This blog post examines how the pandemic has sped up the adoption of telemedicine, enabling remote healthcare delivery and ensuring continuity of care while highlighting challenges related to accessibility and digital literacy. It explores the unprecedented advancements in vaccine technology, showcasing the rapid development and deployment of effective vaccines using both established and novel platforms. The impact on medical education is analyzed, revealing a shift towards online learning and virtual simulations, which have fundamentally altered traditional training methods. The post also addresses the heightened focus on mental health services, emphasizing the increased demand and the need for better preparedness in public health policies. Lastly, it discusses the vulnerabilities exposed in healthcare supply chain management, stressing the importance of building resilient and adaptable systems to meet future demands. Through these insights, the post illustrates how the pandemic has driven significant changes in medicine, shaping its future direction.

##### Background

The COVID-19 pandemic has deeply impacted the field of medicine, driving rapid advancements and necessitating major adaptations across various aspects of healthcare. This global health crisis has spurred change, accelerating the adoption of telemedicine, fostering innovations in vaccine technology, reshaping medical education, emphasizing the importance of mental health services, and revealing weaknesses in healthcare supply chain management.

##### Telemedicine: A Rapid Shift

The pandemic has greatly accelerated the adoption of telemedicine, a technology that allows healthcare providers to deliver care remotely. This shift was driven by the need to minimize face-to-face interactions and ensure the safe delivery of care during the pandemic. Telemedicine has proven particularly beneficial in maintaining the continuity of care, especially in primary care services and mental health care, where in-person visits decreased dramatically. However, concerns about accessibility and digital literacy remain significant challenges.

##### Vaccine Technology: Unprecedented Advancements

The rapid development and deployment of COVID-19 vaccines have been one of the most remarkable achievements during the pandemic. Utilizing both established and new platforms, scientists developed effective vaccines at an unprecedented pace (<u>Peng</u> et al., <u>2021</u>). This effort included the use of nucleic acid-based vaccines, such as mRNA vaccines, which have shown high efficacy and have been rapidly deployed globally (<u>Khoshnood</u> et al., <u>2022</u>). The lessons learned from this rapid development are expected to provide a framework for quick responses to future outbreaks (<u>Hossaini Alhashemi</u> et al., <u>2023</u>).

##### Medical Education: A Major Shift

The pandemic has also had a profound impact on medical education, necessitating rapid adaptations to ensure continuity in training and learning. The closure of medical schools and the shift to online learning and virtual platforms have fundamentally altered how medical curricula are delivered and assessed. This disruption has highlighted the need for sustainable health systems and the importance of preparing medical students for future pandemics (Sharma & Bhaskar, 2020).

##### Mental Health Services: A New Focus

The pandemic has significantly disrupted mental health services, leading to a rise in mental health disorders and emphasizing the importance of mental health as a public health issue. The increased demand for mental health services and the challenges posed by government-imposed restrictions have underscored the need for better preparedness and prioritization of mental health in public health policies (Latoo et al., 2021).

##### Healthcare Supply Chain Management: Building Resilience

The pandemic has exposed significant vulnerabilities in healthcare supply chain management, highlighting the need for resilient and adaptable systems to meet unprecedented demands (Goldschmidt & Stasko, 2022). The failure of traditional supply chain models to respond to the increased demand for critical medical supplies has underscored the importance of strategic and tactical adjustments to ensure the continuous provision of care (Francis, 2020).

Thus, the COVID-19 pandemic has driven significant changes in the field of medicine, highlighting the importance of adaptability, innovation, and resilience in healthcare systems. These changes are likely to have lasting impacts, shaping the future of medicine for years to come.

##### Main text

The COVID-19 pandemic has undeniably reshaped the landscape of medicine, driving rapid advancements and necessitating significant adaptations across various aspects of healthcare. From the adoption of telemedicine to innovations in vaccine technology, changes in medical education, mental health services, and supply chain management, the pandemic has left a lasting impact on the future of medicine.

##### Telemedicine: A New Era of Healthcare Delivery

The COVID-19 pandemic has significantly accelerated the adoption of telemedicine worldwide. The urgent need to minimize face-to-face interactions to curb the spread of the virus led to the rapid implementation of telemedicine services. In Italy, one of the first European countries severely impacted by COVID-19, the healthcare system was overwhelmed, prompting the swift adoption of telemedicine to manage the crisis.

Telemedicine has proven particularly beneficial in maintaining the continuity of care in primary care services, allowing for safe remote triage processes. However, concerns have been raised about its accessibility among older adults, deprived population groups, and individuals with lower digital literacy. In mental health care, the pandemic led to a significant shift from in-person to remote care delivery, with telemedicine usage surging by as much as 6500% in early 2020.

The adoption of telemedicine has also been notable in oncologic care, ensuring the safe delivery of care for patients with thoracic malignancies, although data on its impact on the quality and accessibility of care remain limited. In Portugal, the number of hospital telemedicine consultations increased significantly during the pandemic, highlighting issues such as digital health literacy and technological adoption (Cunha AS et al., 2024; Cunha AS et al., 2024).

In low- and middle-income countries (LMICs), the pandemic has driven the expansion of telemedicine, although its longevity will depend on government regulations, payment policies, and health system redesign. Overall, while telemedicine has proven effective during the pandemic, its future use will require addressing various challenges, including regulatory changes, technological infrastructure, and training for both healthcare professionals and patients (Butaney & Rambhatla, 2022; Cunha AS et al., 2024).

##### Advancements in Vaccine Technology

The COVID-19 pandemic has significantly accelerated advancements in vaccine technology. The rapid development and approval of various vaccine types have been crucial in controlling viral transmission and combating the spread of the disease (Mao et al., 2021).

Researchers have utilized a wide variety of vaccine technologies and platforms, including traditional dead, live, and protein-based vaccines, as well as newer vector-based and gene-based vaccines (Monschein et al., 2021). The intense global effort has led to the rapid development, regulatory approval, and wide-scale distribution of effective vaccines in an impressively short period (Milne et al., 2023).

As of early 2021, over 200 COVID-19 vaccine candidates were in various stages of research and development, covering nearly all existing technologies and platforms (Mao et al., 2021). This rapid development has relied heavily on the application of existing vaccine technologies, ranging from established platforms to emerging technologies with no prior clinical validation (Kiszewski et al., 2021).

The pandemic has also highlighted the need for continuous improvement and adaptation of vaccines to address issues such as long-term protection, side effects, and effectiveness against mutated variants (Abdulla et al., 2021). The development of second-generation vaccines is anticipated to address these challenges and ensure ongoing protection as COVID-19 becomes an endemic disease (Kunkel et al., 2021).

##### Transformations in Medical Education

The COVID-19 pandemic has significantly impacted medical education, necessitating rapid adaptations and innovations to ensure continuity in training and learning. The pandemic led to the closure of medical schools and the barring of patient contact, which disrupted traditional face-to-face teaching and clinical training (Andraous et al., 2022). Consequently, there was a shift towards online learning and virtual teaching methods (Daniel et al., 2021).

Medical students and trainees faced unprecedented challenges, including interruptions in clinical education and increased psychological stress (<u>Akhtar, 2021</u>; <u>Jhajj</u> et al., <u>2022</u>). The quality of education was perceived to have decreased, with 74% of students at McGill University reporting a decline since the start of the pandemic (<u>Jhajj</u> et al., <u>2022</u>).

Despite these challenges, the pandemic also highlighted the need for innovations and technological advancements in medical education. The shift to online platforms and virtual simulations has the potential to change medical education permanently, although the long-term effects are still uncertain (Vaskivska et al., 2021; Nentin et al., 2021). The pandemic has prompted educators to reflect on and potentially reimagine the future models of medical education to better prepare for future crises (Wang W et al., 2024; Sukhera J et al., 2023).

##### Mental Health Services: A New Focus

The COVID-19 pandemic has had a profound impact on mental health services globally. The pandemic has created an unprecedented public health emergency, affecting mental health at political, social, and economic levels (Menculini et al., 2021). Mental health services have faced significant challenges, including disruptions in access and delivery due to government-imposed restrictions and the general burden on healthcare systems (Bauer-Staeb et al., 2021).

The pandemic has led to a “second pandemic” of mental health crises, necessitating health system planning to handle the increased burden (Choi et al., 2020). This includes addressing both existing mental health conditions and new cases of psychological distress emerging from the pandemic (Inchausti et al., 2020).

Mental health services have had to adapt rapidly to these changes. For instance, there has been a need to map early impacts of the pandemic on people with pre-existing mental health conditions and the services they use, as well as to identify strategies to manage these impacts (Sheridan Rains et al., 2021). The pandemic has also highlighted the importance of preparing mental health services for future pandemics (Alexiou et al., 2021).

The impact of the pandemic on mental health has been particularly severe for certain vulnerable groups, who may experience more detrimental effects (Uphoff et al., 2021). The containment measures imposed to control the spread of the virus have had negative consequences on population mental health, leading to an expected increase in both symptomatology and the incidence of mental disorders (Almeda et al., 2021).

##### Supply Chain Management: Building Resilience

The COVID-19 pandemic has significantly impacted supply chain management in healthcare, exposing vulnerabilities and necessitating rapid adaptations to meet unprecedented demands.

The healthcare sector’s supply chain management (SCM) systems faced vulnerabilities due to the intricate nature of COVID-19, fluctuations in demand for specific commodities, and the criticality of dealing with human lives. The pandemic created global health and economic disruptions, with hospitals and other healthcare providers being hit particularly hard (<u>Francis,</u> <u>2020</u>).

During the early months of the pandemic, tertiary multihospital systems were highly viewed as the go-to hospitals for handling rapid healthcare challenges caused by the increasing number of COVID-19 cases (Alswailem et al., 2021). The traditional cost-effective supply chain paradigm failed to respond to the increased demand, putting healthcare workers at a much higher infection risk relative to the general population (Baloch G et al., 2023).

The pandemic exposed significant fragilities in the configuration of global healthcare supply chains, felt acutely by citizens, patients, and healthcare workers across Canada (<u>Snowdon</u> et al., <u>2022</u>). Provincial healthcare supply chain teams worked to rapidly stabilize their supply chains, indicating emerging features of healthcare supply chain resilience (<u>Snowdon</u> et al., <u>2022</u>).

The failure of the global supply chain became rapidly apparent at the beginning of the pandemic when healthcare organizations were left without supplies needed to care for patients, indirectly impacting patient morbidity and mortality (Goldschmidt & Stasko, 2022). The pandemic has shown the importance of resilience in the pharmaceutical industry to deal with unexpected disruptions and high demand from patients and authorities (Belghith M & Bouajaja S, 2024).

Infection prevention and control practices remain the foundation of healthcare-associated infection prevention and outbreak control efforts. However, issues in supply chain management can hinder these efforts, as exemplified by various public health emergencies (<u>Mehrotra P & Desai A, 2024</u>). The pandemic has highlighted the many challenges that provincial health systems have experienced while scaling health services to protect Canadians from viral transmission and support care for those who get infected (<u>Snowdon</u> et al., <u>2021</u>).

The COVID-19 pandemic has underscored the critical need for resilient and adaptable supply chain management in healthcare to ensure the continuous provision of essential medical supplies and to safeguard public health.

#### Conclusion

The COVID-19 pandemic has significantly reshaped the future of medicine, driving rapid advancements and necessitating major adaptations across various aspects of healthcare. Telemedicine emerged as an essential tool, ensuring continuity of care while minimizing physical interactions, although challenges in accessibility and digital literacy remain. Vaccine technology saw unprecedented acceleration, with rapid development and deployment of various vaccine platforms, setting a new standard for future pandemics. Medical education underwent a major shift towards online learning, highlighting the need for innovative and flexible training methods. Mental health services faced immense pressure, prompting a reevaluation of mental health care delivery and preparedness for future crises. Lastly, the pandemic exposed weaknesses in healthcare supply chains, emphasizing the importance of building resilient and adaptable systems. Collectively, these changes represent a transformative period in medicine, with lasting effects on healthcare delivery, education, and infrastructure, paving the way for a more resilient and responsive healthcare system in the future.

##### References

Abdulla, Al-Bashir, Al-Salih, Aldamen & Abdulazeez A Summary of the SARS-CoV-2 Vaccines and Technologies Available or under Development. Pathogens (Basel, Switzerland) (2021).

Akhtar Exploring the Influence of a Pandemic on Medical Education. Missouri medicine (2021).

Alexiou, et al. A Survey of Psychiatric Healthcare Workers’ Perception of Working Environment and Possibility to Recover Before and After the First Wave of COVID-19 in Sweden. Frontiers in psychiatry (2021).

Almeda, García-Alonso & Salvador-Carulla Mental health planning at a very early stage of the COVID-19 crisis: a systematic review of online international strategies and recommendations. BMC psychiatry (2021).

Alswailem, et al. COVID-19 Intelligence-Driven Operational Response Platform: Experience of a Large Tertiary Multihospital System in the Middle East. Diagnostics (Basel, Switzerland) (2021).

Andraous, Amin & Allam The “new normal” for medical education during and post-COVID-19. Education for health (Abingdon, England) (2022).

Antonacci G et al. Healthcare professional and manager perceptions on drivers, benefits, and challenges of telemedicine: results from a cross-sectional survey in the Italian NHS. BMC health services research (2023).

Baloch G, Gzara F & Elhedhli S Risk-based allocation of COVID-19 personal protective equipment under supply shortages. European journal of operational research (2023).

Bauer-Staeb et al. The early impact of COVID-19 on primary care psychological therapy services: A descriptive time series of electronic healthcare records. EClinicalMedicine (2021).

Belghith M & Bouajaja S [Implementation of an Advanced Planning and Scheduling System in a pharmaceutical context]. Annales pharmaceutiques francaises (2024).

Bokolo Exploring the adoption of telemedicine and virtual software for care of outpatients during and after COVID-19 pandemic. Irish journal of medical science (2021).

Butaney & Rambhatla The impact of COVID-19 on urology office visits and adoption of telemedicine services. Current opinion in urology (2022).

Choi, Heilemann, Fauer & Mead A Second Pandemic: Mental Health Spillover From the Novel Coronavirus (COVID-19). Journal of the American Psychiatric Nurses Association (2020).

Cunha AS, Pedro AR & V Cordeiro J Challenges of Using Telemedicine in Hospital Specialty Consultations during the COVID-19 Pandemic in Portugal According to a Panel of Experts. Acta medica portuguesa (2024).

Daniel, et al. An update on developments in medical education in response to the COVID-19 pandemic: A BEME scoping review: BEME Guide No. 64. Medical teacher (2021).

Engelbrecht M, Ngqangashe Y, Mduzana L, Sherry K & Ned L Disability inclusion in African health systems’ responses during COVID-19: A scoping review. African journal of disability (2023).

Francis COVID-19: Implications for Supply Chain Management. Frontiers of health services management (2020).

Goldschmidt & Stasko The downstream effects of the COVID-19 pandemic: The supply chain failure, a wicked problem. Journal of pediatric nursing (2022).

Hoffer-Hawlik, et al. Leveraging Telemedicine for Chronic Disease Management in Low- and Middle-Income Countries During Covid-19. Global heart (2020).

Hossaini Alhashemi, Ahmadi & Dehshahri Lessons learned from COVID-19 pandemic: Vaccine platform is a key player. Process biochemistry (Barking, London, England) (2023).

Inchausti, MacBeth, Hasson-Ohayon & Dimaggio Psychological Intervention and COVID-19: What We Know So Far and What We Can Do. Journal of contemporary psychotherapy (2020).

Jhajj, et al. Impact of Covid-19 on Medical Students around the Globe. Journal of community hospital internal medicine perspectives (2022).

Khoshnood, et al. Viral vector and nucleic acid vaccines against COVID-19: A narrative review. Frontiers in microbiology (2022).

Kiszewski, Cleary, Jackson & Ledley NIH funding for vaccine readiness before the COVID-19 pandemic. Vaccine (2021).

Kunkel, Madani, White, Verardi & Tarakanova Modeling coronavirus spike protein dynamics: implications for immunogenicity and immune escape. Biophysical journal (2021).

Latoo, et al. The COVID-19 pandemic: an opportunity to make mental health a higher public health priority. BJPsych open (2021).

Lin YK et al. Global prevalence of anxiety and depression among medical students during the COVID-19 pandemic: a systematic review and meta-analysis. BMC psychology (2024).

Maluleke K et al. Co-creation of a novel approach for improving supply chain management for SARS-CoV-2 point of care diagnostic services in Mopani District, Limpopo Province: nominal group technique. Frontiers in public health (2024).

Mao, et al. COVID-19 vaccines: progress and understanding on quality control and evaluation. Signal transduction and targeted therapy (2021).

Mehrotra P & Desai A Resource sustainability and challenges in the supply chain: implications for infection prevention. Current opinion in infectious diseases (2024).

Menculini, et al. Access to Mental Health Care during the First Wave of the COVID-19 Pandemic in Italy: Results from the COMET Multicentric Study. Brain sciences (2021).

Milne et al. Independent control of COVID-19 vaccines by EU Official Control Authority Batch Release: challenges, strengths and successes. NPJ vaccines (2023).

Monschein, et al. [The corona pandemic and multiple sclerosis: vaccinations and their implications for patients-Part 2: vaccine technologies]. Der Nervenarzt (2021).

Nentin, Gabbur & Katz A Shift in Medical Education During the COVID-19 Pandemic. Advances in experimental medicine and biology (2021).

Nimgaonkar, et al. Impact of telemedicine adoption on accessibility and time to treatment in patients with thoracic malignancies during the COVID-19 pandemic. BMC cancer (2021).

Olszewska B, Zaryczańska A, Nowicki RJ & Sokołowska-Wojdyło M Rare COVID-19 vaccine side effects got lost in the shuffle. Primary cutaneous lymphomas following COVID-19 vaccination: a systematic review. Frontiers in medicine (2024).

Peng, et al. Advances in the design and development of SARS-CoV-2 vaccines. Military Medical Research (2021).

Pulido-Fuentes et al. Internal dynamics within primary care teams in two Spanish regions during the COVID-19 pandemic: a qualitative study. BMC primary care (2022).

Rahman MW, Hasan MM, Palash MS & Asaduzzaman M Medical education in Bangladesh from Student and Teacher’s Perspective: Impact and challenges of the COVID-19 pandemic. MedEdPublish (2016) (2024).

Sharma & Bhaskar Addressing the Covid-19 Burden on Medical Education and Training: The Role of Telemedicine and Tele-Education During and Beyond the Pandemic. Frontiers in public health (2020).

Sheridan Rains et al. Early impacts of the COVID-19 pandemic on mental health care and on people with mental health conditions: framework synthesis of international experiences and responses. Social psychiatry and psychiatric epidemiology (2021).

Snowdon, Saunders & Wright Key Characteristics of a Fragile Healthcare Supply Chain: Learning from a Pandemic. Healthcare quarterly (Toronto, Ont.) (2021).

Snowdon, Saunders & Wright The Emerging Features of Healthcare Supply Chain Resilience: Learning from a Pandemic. Healthcare quarterly (Toronto, Ont.) (2022).

Sukhera J, et al. Digging Deeper, Zooming Out: Reimagining Legacies in Medical Education. Academic medicine : journal of the Association of American Medical Colleges (2023).

Tsoli F et al. Difficult-to-Engage Patients with Severe Mental Illness in Rural Community Settings: Results of the Greek Hybrid Assertive Community Treatment Model of Mental Healthcare. Journal of clinical medicine (2024).

Uphoff, et al. Mental health among healthcare workers and other vulnerable groups during the COVID-19 pandemic and other coronavirus outbreaks: A rapid systematic review. PloS one (2021).

Vaskivska, Palamar, Kravtsova & Khodakivska TRANSFORMATION OF THE LEARNING PROCESS IN HIGHER EDUCATION INSTITUTIONS UNDER THE INFLUENCE OF THE PANDEMIC COVID-19. Wiadomosci lekarskie (Warsaw, Poland : 1960) (2021).

Wang W, Li G & Lei J The impact of COVID-19 on medical students. GMS journal for medical education (2024).

Wilczewski, et al. Providers’ Perspectives on Telemental Health Usage After the COVID-19 Pandemic: Retrospective Analysis. JMIR formative research (2022).

## Example 5: Press release

User’s selected inputs:

1. Title and abstract from the publication by Kuznetsov et al. [37];
2. Organization - Insilico Medicine

#### Insilico Medicine Introduces COSMIC: A Game-Changer in Fast and Accurate Molecular Modeling

##### Main text

Scientists at Insilico Medicine have developed COSMIC (COnformation Space Modeling in Internal Coordinates), an innovative adversarial generative framework that significantly enhances the speed and accuracy of molecular conformation modeling. Published in the Journal of Chemical Information and Modeling, this research offers a promising advancement in ligand and structure-based drug discovery.

##### Tackling a Major Challenge in Drug Discovery

Understanding the 3D arrangements of molecules, known as conformations, is vital in the realm of drug discovery. These structures influence a molecule’s docking poses and binding energy, crucial for understanding interactions with biological targets. Traditional methods, while accurate, come with high computational costs and slow sampling speeds, creating a bottleneck in the drug discovery process. COSMIC addresses these challenges by providing a faster, more efficient way to generate molecular conformations without compromising on quality (<u>Kuznetsov M</u> et al., <u>2024</u>).

##### Novel Two-Stage Generative Process

COSMIC operates through an innovative two-stage generative process. Initially, it constructs a molecular conformation in a rotation and translation-invariant representation known as internal coordinates. This is followed by predicting the distances between neighboring atoms and performing a few rapid optimization steps to refine the initial conformation. This method ensures that the generated conformations are both physically plausible and energetically favorable.

##### Key Advancements and Impact

The COSMIC framework demonstrates a significant improvement in the speed and accuracy of molecular conformation generation. Unlike previous models, COSMIC introduces a new energy-based metric called RED (Relative Energy Difference) to ensure the physical plausibility of generated conformations. This metric allows for large-scale quantitative comparisons of energetic performance.

The study’s extensive experiments and quantitative comparisons with existing models show that COSMIC outperforms traditional methods and non-diffusion neural baselines in several key metrics, including space coverage and distribution matching. The framework’s ability to generate diverse and physically plausible conformations quickly positions it as a significant advancement in the field.

##### Future Directions and Potential Applications

The implications of this research are far-reaching. By providing a robust and efficient method for molecular conformation generation, COSMIC has the potential to accelerate the drug discovery process, making it easier to identify promising drug candidates and understand their interactions with biological targets. This could lead to the development of new therapies for a range of diseases, from cancer to autoimmune disorders.

Future research will likely focus on scaling the COSMIC framework to handle even more complex molecules and integrating it with other computational tools to provide a comprehensive solution for drug discovery.

*“In the long-term, we envision COSMIC being integrated into various computational platforms, enhancing the capabilities of researchers worldwide,” adds Dr. Zhavoronkov.*

#### Conclusion

Insilico Medicine’s COSMIC framework represents a significant leap forward in molecular conformation modeling. By combining speed, accuracy, and physical plausibility, COSMIC stands to revolutionize the way researchers approach drug discovery, paving the way for new treatments and therapies. As the field continues to evolve, the contributions of COSMIC will undoubtedly play a crucial role in shaping the future of computational chemistry and molecular biology.

##### References

Kuznetsov M et al. COSMIC: Molecular Conformation Space Modeling in Internal Coordinates with an Adversarial Framework. Journal of chemical information and modeling (2024).

## Validation

With the increased usage of LLMs, the necessity for robust methods to benchmark LLM-generated text has significantly increased. Currently proposed methods and benchmarks involve assessing coherence, factual accuracy, fluency, and alignment with human values through diverse benchmarks and metrics [38], [39]. Advanced evaluation techniques are also being explored, incorporating fine-grained criteria such as lexical diversity and sentence complexity [40]. Trends in this area suggest a growing interest in multi-scenario meta-evaluation benchmarks and the use of LLMs themselves for evaluation purposes, although human oversight remains crucial to capture complex errors [41], [42].

Despite these advancements, there is a notable lack of universally approved methodologies for evaluating LLM-generated texts, particularly complex ones such as scientific articles. To address this gap, we aimed to develop our own validation system for analyzing generated texts. We began with extensive analysis of scientific documents to identify the key metrics indicative of high-quality text that resulted in identification of 8 scores for future grading (Supplementary Table 2). Subsequently, we manually devised a grading system that delineates criteria for assigning scores ranging from 0 (lowest) to 10 (highest) for each of these metrics. Each of the eight scoring systems was then integrated into a prompt for GPT-4, instructing the model to evaluate the given text based on the specified criteria.

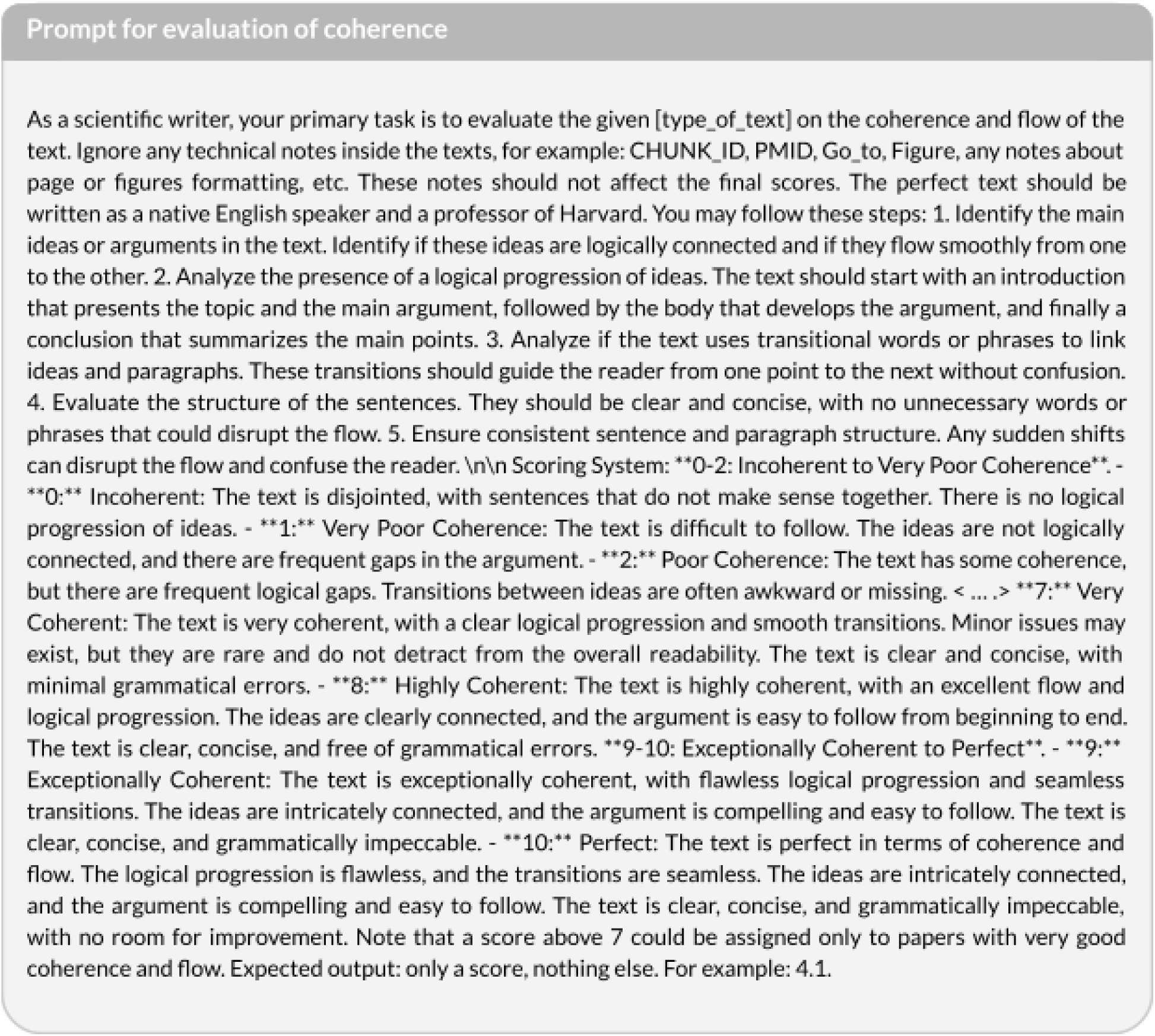

To assess the quality of writing produced by several AI writing assistants, we generated ten review articles using DORA and 3 other AI writing assistants out of the most popular enterprise solutions. The topics for these articles were selected from actual research published in Nature Reviews. Selected Nature papers served as positive controls, representing the text that should be assigned with the highest possible quality scores. Additionally, we created paired negative controls: fabricated papers designed to receive the lowest possible scores, generated using GPT-4 with manual corrections.

To evaluate the first 7 criteria (Paper structure, Relevance to topic, Writing quality, Coherence and flow, Depth and detail, Tone and style, Information density) the citations were excluded from the text to avoid bias, as citation formats and placements varied across sources. Each text was then assessed on each criterion individually using corresponding prompts for the GPT-4 model. Initial scores ranged from 0 to 10 and were normalized such that the minimum score corresponded to the negative control and the maximum score to the Nature review article. These normalized scores were then scaled from 0 to 100. For citation quality, citations were selected randomly from each text and manually evaluated for their relevance to the corresponding facts mentioned. The grading system ranged from 0 for hallucinated citations to 5 for perfect correspondence. These scores were also normalized to a 0 to 100 scale to align with the other evaluation criteria.

First, the analysis has shown that real papers consistently show great performance that generated papers that could testify to the ability of LLM to capture the difference in generated text using identified criteria. Furthermore, the results for DORA consistently rank high across most criteria, including Writing Quality, Coherence and Flow, and Tone and Style, comparable to those of high-quality review papers (Fig. 4). Among the AI writers analyzed, DORA’s performance is notably superior. The only criterion where DORA scores lower is Information Density. However, our manual analysis indicates that this might be due to DORA’s inclusion of connective sentences, as well as analysis and interpretation of facts. Given that DORA’s Depth and Detail score, which measures the richness of information, is high, the lower Information Density should not be viewed as a disadvantage, in contrast it might reflect DORA’s design to integrate analysis and interpretation, thereby enhancing readability. This should be further studied in subsequent work.

## Conclusion

Our work has demonstrated that DORA represents a significant advancement in the realm of AI-driven research and manuscript generation. By leveraging LLM agents and a meticulously designed pipeline, DORA simplifies the drafting process while maintaining high flexibility and user-specific customization. This innovative approach allows researchers to produce high-quality, coherent documents with minimal effort, thereby accelerating the scientific research process. The integration of Insilico Medicine’s advanced platforms, such as PandaOmics and Precious3GPT, further enhances DORA’s capabilities, enabling it to autonomously gather, synthesize information, and substantiate scientific hypotheses with robust data. The incorporation of a shared memory system and specific order of sections generator ensures logical flow and coherence across all sections of the document, making DORA not just a tool for generating research publications, but a catalyst for scientific innovation.

We envision the integration of additional AI tools that can retrieve data from a broader range of sources, both internal and external. Plans are in place to expand the range of templates to cover more scientific disciplines and document types, including templates tailored for various scientific journals. This expansion will allow DORA to cater to a wider audience and address diverse research needs. Furthermore, user feedback will be continuously incorporated to refine and enhance DORA’s functionalities, ensuring that it remains at the forefront of AI-driven research tools.

However, it is important to acknowledge the limitations and challenges associated with DORA. While the reliance on predefined templates and LLMs offers significant advantages, it also presents certain technical limitations. For instance, the quality of the generated content is highly dependent on the quality of the prompt provided, custom input data, and the specificity of the templates. Additionally, the use of LLMs can sometimes result in the generation of content that lacks depth or contains inaccuracies. To mitigate these challenges, we have implemented strategies such as the integration of RAG systems, which reduce the risk of hallucinations and ensure transparency by providing relevant citations. Furthermore, the shared memory system enhances the logical flow and coherence of the final document, ensuring that major results and insights are consistently discussed across all sections. Despite these measures, continuous monitoring and refinement are necessary to address any emerging issues and maintain the high quality of the generated content. Humans are an essential part of the process and they should verify the output of tools like DORA before having them released publically or further acted upon.

In summary, DORA stands as a groundbreaking tool in the field of AI-driven research publication generation. Its unique features and capabilities offer significant advantages over traditional methods, streamlining the drafting process and empowering researchers to focus on their core scientific endeavors. While there are limitations and challenges to be addressed, the future prospects of DORA are bright, with ongoing advancements and user feedback driving continuous improvement. As AI continues to shape the landscape of academic science, tools like Science42: DORA will remain indispensable in accelerating literature review, and translating complex results into structured and accurate content.

### Future perspectives

The development of DORA represents a significant step in the automation of scientific writing, creating the way for an even more ambitious vision: the complete automation of scientific research processes. Looking ahead, our vision for DORA’s evolution is to create an environment where scientific research can be conducted entirely autonomously, requiring only an objective-prompt from the user. This transformation would see DORA evolve from an AI-driven writing assistant into a comprehensive autonomous research system, capable of designing experiments, analyzing data, and generating results with minimal human intervention. Achieving this would involve enhancing the current multi-agent system with advanced prompt techniques, such as self-reflection loops, integrating multiple novel tools, and completing integration with the Robotics Lab. This scenario could significantly accelerate the research process, reduce human error, and allow scientists to focus on higher-level conceptual work and innovation. This future vision underscores the transformative potential of DORA in reshaping the landscape of scientific research and communication.

### Materials availability

DORA is available for free during the trial period at https://insilico.com/science42/dora.

### Note on AI usage for publication drafting

DORA was used to prepare the draft of the current manuscript. The initially generated draft was manually curated and updated to .

## Acknowledgements

We thank Ms. Elizaveta Ekimova for her technical assistance with figure design.

## Supplementary data

**Supplementary Table 1.**
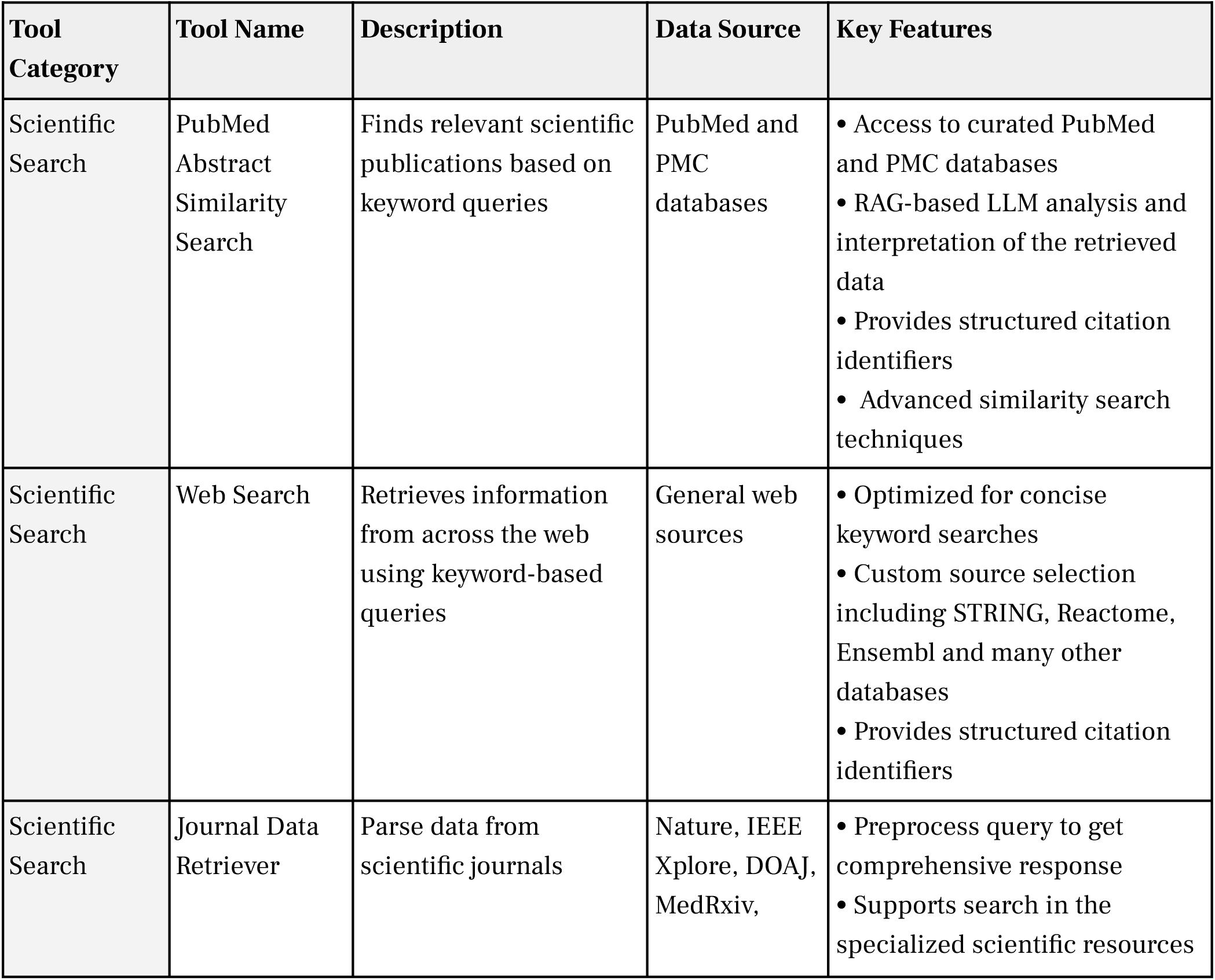
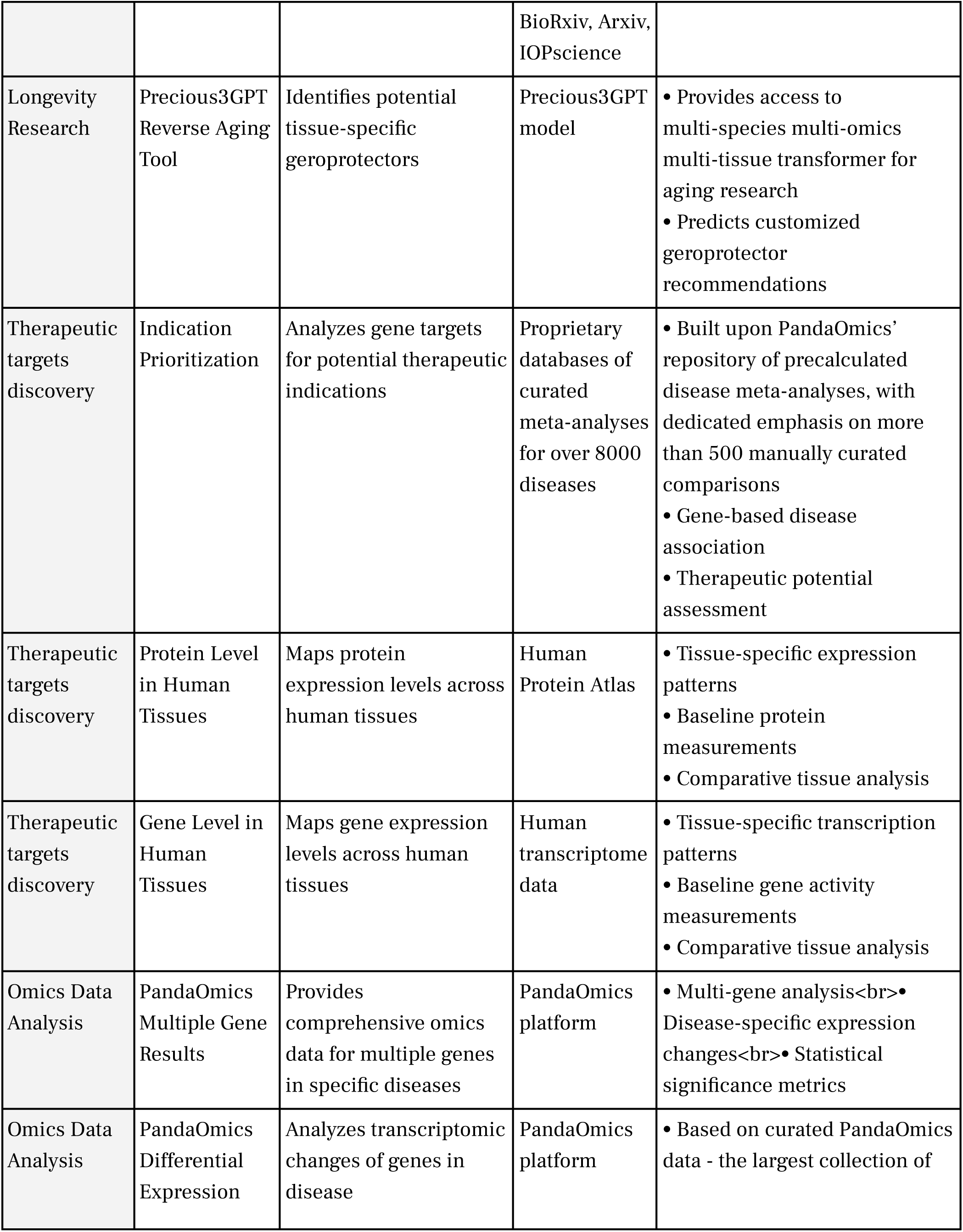
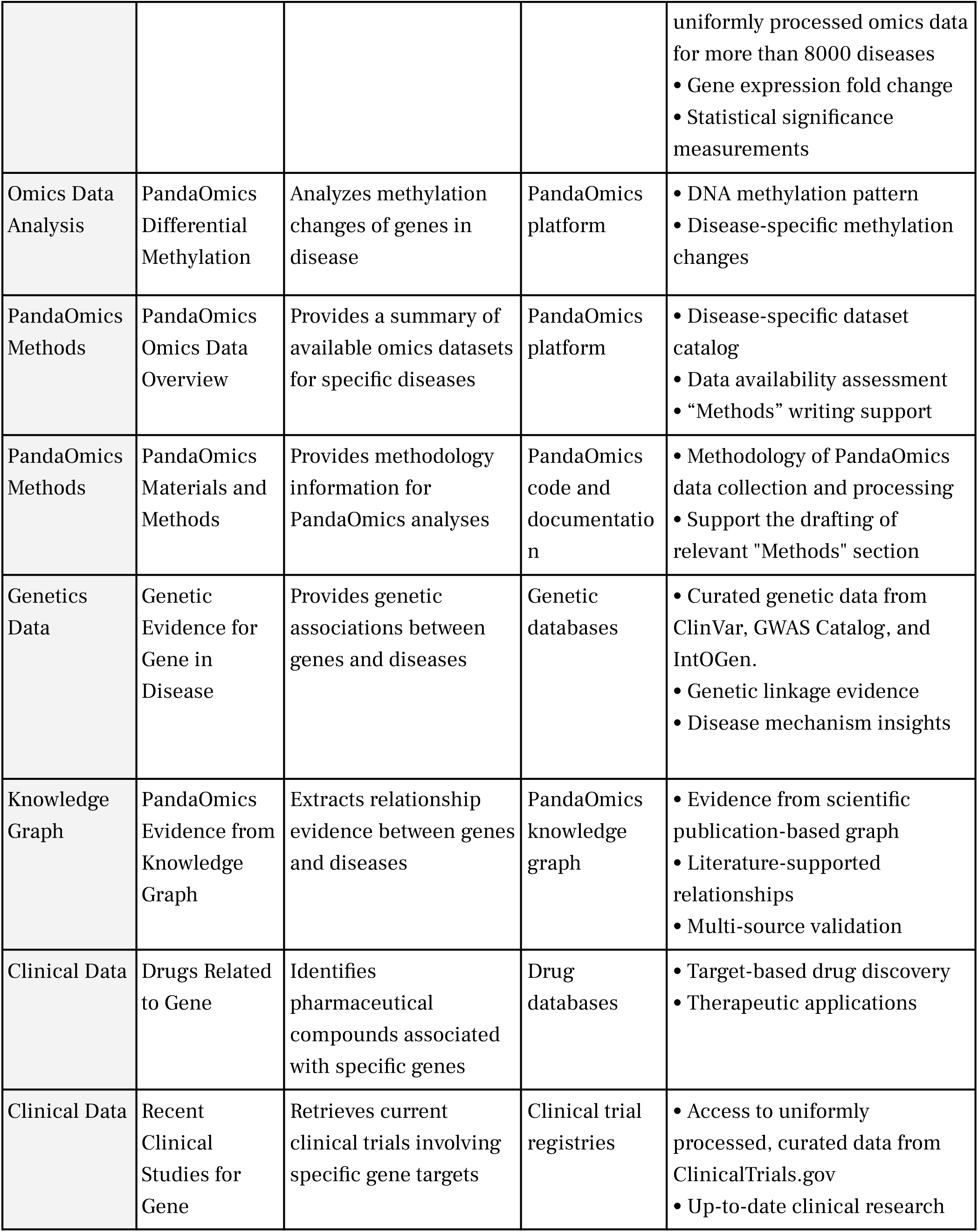
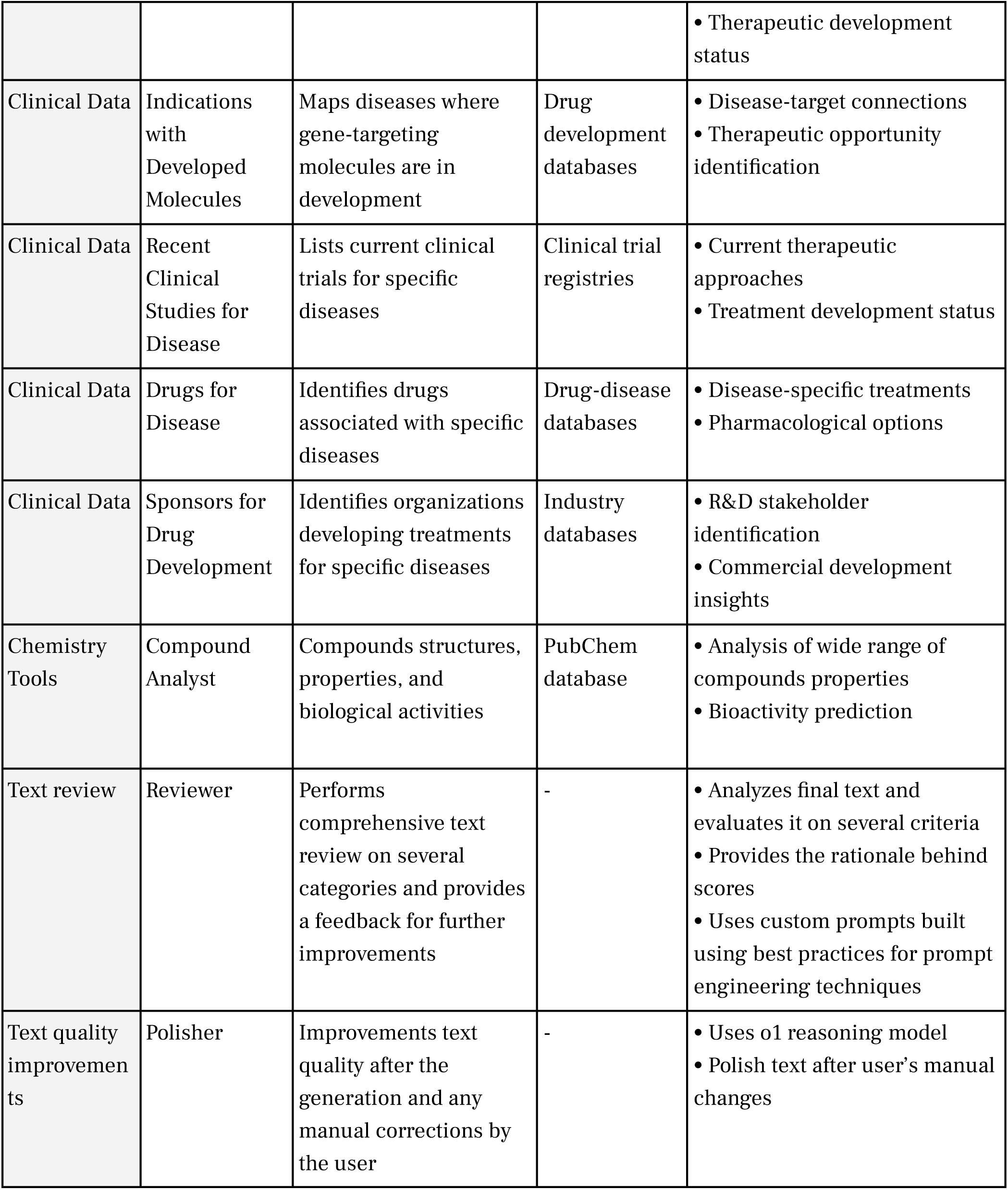
The list of tools available in DORA.

**Supplementary Table 2.** Criteria used for validation prompts construction.

|  | Name | Description |
| --- | --- | --- |
| 1 | Paper structure:<br>Organized and logical layout suitable for the selected document type | Evaluates the logical organization and coherence of the text, focusing on the clear division of sections, smooth flow of ideas, and the effectiveness of the conclusion. To achieve a high score, the text must exhibit well-defined sections, seamless transitions between ideas, a concise and comprehensive conclusion, and a structure that is highly suitable for its type. |
| 2 | Relevance to topic:<br>directly addresses the subject matter. | Evaluates how closely the content aligns with the main topic and subtopics of the text. To achieve a high score, ensure the text provides a comprehensive, in-depth discussion of the main topic, integrates diverse perspectives, and supports conclusions with high-quality, relevant information. |
| 3 | Writing quality: high standard of grammar and style | This criterion evaluates the grammatical correctness, punctuation accuracy, and appropriateness of scientific language in the text. To achieve a high score, the text must exhibit flawless grammar, precise punctuation, and clear, appropriate scientific language. |
| 4 | Coherence and flow: smooth and logical progression of the idea | This criterion evaluates the logical progression and smoothness of the text. To achieve a high score, ensure that the main ideas are logically connected, transitions between ideas are seamless, and the text flows naturally from introduction to conclusion. Sentences and paragraphs should be clear, concise, and consistently structured to maintain readability. |
| 5 | Depth and detail: comprehensive and thorough analysis | Evaluates the thoroughness and specificity with which the topic is explored. To achieve a high score, the text must cover all relevant aspects comprehensively, present highly complex ideas, and provide well-supported arguments with specific examples and clear explanations. |
| 6 | Tone and style: appropriate and engaging language | Evaluates the clarity, coherence, and sophistication of the writing. To achieve a high score, the text must maintain a professional and academic tone, be free from bias, and demonstrate a high level of linguistic and rhetorical skill. |
| 7 | Information density: enrichment with valuable content and facts | This criterion evaluates the concentration of relevant, unique scientific information in the main section of the text relative to its length. A high score is achieved by presenting a concise, varied array of scientific facts without unnecessary repetition, filler content, or off-topic information. |
| 8 | Citation quality: accuracy and relevance in citations and associated | Citation quality assesses the accuracy and relevance of citations and their metadata, with high scores awarded for precise alignment between the cited fact in the text and the original source, ensuring scholarly integrity and reliability. |
|  | metadata |  |

